# Backtracking of influenza polymerase upon consecutive incorporation of nucleoside analogue T1106 directly observed by high-resolution cryo-electron microscopy

**DOI:** 10.1101/2022.06.10.495428

**Authors:** Tomas Kouba, Anna Dubankova, Petra Drncova, Elisa Donati, Pietro Vidossich, Valentina Speranzini, Alex Pflug, Johanna Huchting, Chris Meier, Marco De Vivo, Stephen Cusack

**Affiliations:** European Molecular Biology Laboratory, 71 Avenue des Martyrs, CS 90181, 38042 Grenoble Cedex 9, France; Organic Chemistry, Department of Chemistry, Hamburg University, Martin-Luther-King-Platz 6, D-20146 Hamburg, Germany; Molecular Modeling & Drug Discovery Lab, Istituto Italiano di Tecnologia, Via Morego 30 - Genova, 16163, Italy

**Keywords:** Influenza, RNA-dependent RNA polymerase, antiviral drug, nucleoside analogue, T705 (favipiravir), T1106, cap-dependent transcription, backtracking, single-particle cryo-electron microscopy, molecular dynamics

## Abstract

The broad-spectrum antiviral pseudobase T705, a fluorinated pyrazinecarboxamide, is incorporated via its triphosphate form into nascent viral RNA by viral RNA-dependent RNA polymerases. Since it mimics guanine or adenine it can act as a mutagen, whereas consecutive incorporation leads to chain termination. Here we examine the structural basis for incorporation and stalling for the case of influenza polymerase, using T1106-TP, the nucleotide form of T1105, the de-fluoro analogue of T705. We used a specially designed template that allows single T1106-MP incorporation at a defined site followed by consecutive T1106-MP incorporation and stalling four nucleotides later, as demonstrated by biochemical analysis. A high-resolution cryoEM structure of influenza A/H7N9 polymerase, stalled after transcribing this template, revealed that the entire product-template duplex has backtracked by five nucleotides. Consequently, the singly incorporated T1106-MP resides at the +1 position and forms an unexpected wobble base-pair with a U in the template. The relative stability of the canonical and wobble T1106:U base-pairs in different contexts is investigated by molecular dynamics simulations. Using a different template and influenza B polymerase we also observe stalling after double incorporation of T1106-MP and structural analysis showed again that backtracking occurs, this time by four nucleotides. These results show that, at least in early elongation, consecutive T1106-MP incorporation into the product destabilises the proximal end of the product-template duplex, promoting irreversible backtracking until a more favourable overall configuration is achieved. These results give new insight into the unusual mechanism of chain termination by pyrazinecarboxamide base analogues.

## Introduction

Many concerning human pathogens are RNA viruses and, except for retroviruses, all encode an RNA-dependent RNA polymerase that is responsible for replication and transcription of the RNA genome. The core activity of this enzyme is template dependent nucleotide addition to a growing product RNA and this is catalysed by a conserved active site formed by several characteristic structural motifs. Nucleoside analogues that impede RNA synthesis by various mechanisms therefore have the potential to be broad-spectrum inhibitors of RNA virus replication. One such inhibitor is T705 (favipiravir), a fluorinated pyrazinecarboxamide pseudo base, originally developed against influenza virus (Furuta *et al*, 2002; Shiraki & Daikoku, 2020; Sidwell *et al*, 2007), but also active against several other viruses (Lagocka *et al*, 2021) including SARS-CoV-2 (Kaptein *et al*, 2020; Naydenova *et al*, 2021; Peng *et al*, 2021; Shannon *et al*, 2020). T705 is converted by cellular enzymes through ribosylation and phosphorylation into T705 triphosphate (T705-TP), which is incorporated by the viral polymerase opposite either U or C in the template, due to the ambiguous base-pairing endowed by the rotatable carboxamide group of the pseudobase (Jin *et al*, 2013). It has been reported that ATP and GTP are favoured over T705-TP as substrates by discriminatory factors of respectively 30-fold and 19-fold (Jin *et al*., 2013). Two mechanisms of viral inhibition by T705 have been proposed, lethal mutagenesis and chain termination. Firstly, dispersed individual incorporation of T705-MP into the RNA product does not lead to chain termination, but generates a product RNA that has ambiguous coding specificity, consistent with an enrichment in G to A and C to U mutations being observed in viral RNAs upon treatment with T705 (Baranovich *et al*, 2013; Goldhill *et al*, 2019). Secondly, it has been observed that consecutive incorporation of T705-MP into product RNA leads to chain-termination consistent with complete inhibition of RNA synthesis being observed at high T705 concentrations (Jin *et al*., 2013; Wang *et al*, 2021). Although no favipiravir-resistant viruses have appeared in clinical trials (Takashita *et al*, 2016), recently it has been possible to select for an influenza polymerase mutant that has low susceptibility to T705. The PB1 K229R mutation in the context of the 2009 pH1N1 strain shows reduced incorporation of T705, probably due to enhanced fidelity, but also reduced polymerase activity that could be compensated by an accompanying PA P653L mutation (Goldhill *et al*, 2018). It has since been shown that an alternative, naturally occurring mutation (PA N321K) can partially restore polymerase activity to the PB1 K229R resistant virus in the context of a more recent H1N1 strain, suggesting that under certain circumstances it might be possible that T705 resistance could be conferred by a single mutation (Goldhill *et al*, 2021).

Here we investigate the structural basis for incorporation and stalling of influenza polymerase by the active triphosphate form of nucleoside-analogue T1106, derived from nucleobase T1105, the de-fluoro analogue of T705 (Furuta *et al*, 2013). This compound is more stable, more readily metabolised and more potent than T705 (Barauskas *et al*, 2017; Huchting *et al*, 2019; Huchting *et al*, 2018; Huchting *et al*, 2017). For instance, using purified influenza vRNPs, the IC50 of T1106-TP (hereafter designated as simply T1106) was found to be 0.48 and 0.69 μM for influenza A and B respectively (Huchting *et al*., 2018). Since T1106 also substitutes for ATP and GTP, we designed combinations of capped primer and non-native template sequences that only require incorporation of UTP and CTP in the early steps of cap-dependent transcription, but depend on T1106 for insertion opposite specifically placed U (single incorporation) or UC (double incorporation) sites in the template. Using recombinant polymerase from influenza A/Zhejiang/DTID-ZJU01/2013(H7N9) (Chen *et al*, 2013), subsequently referred to as A/H7N9 polymerase we find biochemically that RNA synthesis is indeed stalled upon double incorporation. A cryoEM structure at 2.51 Å of the stalled complex was determined that showed that the entire template-product duplex had backtracked by five nucleotides. This unusual structure is compared with a 2.48 Å structure of a normal early elongation complex of the same A/H7N9 polymerase, in which no T1106 was used and the transcription reaction was stalled, after incorporation of 5 nucleotides, by addition of the non-hydrolysable CTP analogue, cytidine-5’-[(α,β)-methyleno]triphosphate (CMPcPP). Since these are the first high-resolution structures of a complete A/H7N9 polymerase, particular features of this polymerase will be highlighted, including the cryoEM structures of the homodimeric apo-and 5′ hook bound forms. To confirm that backtracking is a general phenomenon resulting from double incorporation of T1106 in early transcription, we obtained a 2.58 Å structure of stalled influenza B polymerase, backtracked by 4 nucleotides, using a slightly different template that only has a double, and no single, T1106 incorporation site. This backtracked structure can be compared with a previously determined structure of a normal early elongation complex of the same polymerase (PDB:6QCT, 3.2 Å) (Kouba *et al*, 2019).

This is the first direct structural proof that nucleotide analogue incorporation can cause influenza polymerase to backtrack, although T1106 induced backtracking has been detected by single molecule techniques for positive-strand RNA viral polymerases (Dulin *et al*, 2017; Janissen *et al*, 2021)(Seifert *et al*., https://www.biorxiv.org/content/10.1101/2020.08.06.240325v2). Backtracking is used by cellular (Cheung & Cramer, 2011), bacterial (Abdelkareem *et al*, 2019) and some viral (Malone *et al*, 2021) polymerases to proofread and potentially correct misincorporation, provided a means of excising the backtracked nucleotides is available.

## Results

### Biochemical analysis of influenza polymerase transcription with and without T1106 incorporation

In this study, we have made the first use of full-length, recombinant heterotrimeric polymerase from A/Zhejiang/DTID-ZJU01/2013(H7N9), although crystal structures of a truncated construct have been described previously (Wandzik *et al*, 2020). This is a human isolate of an avian strain possessing adaptive mutations PB2/E627 and N701 (Chen *et al*., 2013). In high-salt buffer, the full-length A/H7N9 polymerase purifies primarily as dimers of heterotrimers (Figure S1A,C,D), as previously observed for the human A/H3N2 and avian A/H5N1 polymerases (Fan *et al*, 2019). CryoEM structures of the apo- and 5′ hook-bound dimeric forms are described below. Upon incubation with the full vRNA promoter in low-salt buffer, a large fraction of the promoter bound complex is monomeric (Figure S1B, D).

To obtain a stalled early elongation complex of the A/H7N9 polymerase we performed a transcription reaction using the previously described extended 3’ end 18+3-mer template (Kouba *et al*., 2019), capped 13-mer primer and 14-mer 5’ vRNA activator ‘hook’, together with ATP and GTP and non-hydrolysable CMPcPP (cytidine-5’-[(α,β)-methyleno]triphosphate)(Figure 1A). The template translocated by five nucleotides from the initiation state before stalling at the first guanine in the template sequence at position (9) (bracketed numbers indicate the position from the native 3’ end of the template). At this point, the principle product is a capped 18-mer, as confirmed by the unambiguous base pair identity of the product-template duplex visualised in the cryoEM density at 2.48 Å resolution (Figure S2A).

**Figure 1.**
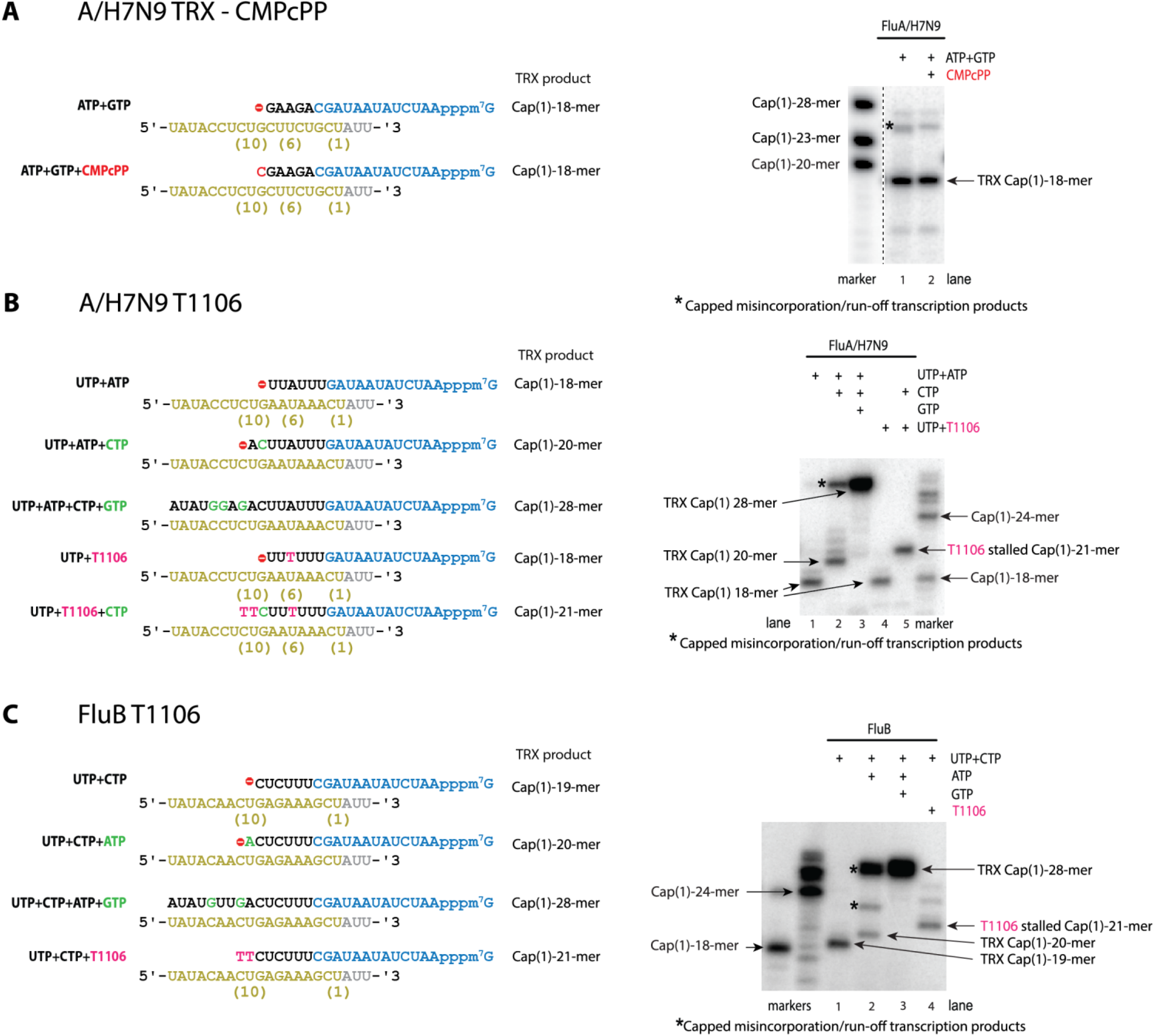
Templates and transcription (TRX) products. **A.(Left)** 18+3-mer template (yellow) used for A/H7N9 transcription reactions with capped 13-mer primer (blue) and ATP, GTP (black) with and without CMPcPP (red). **(Right)** Gel showing elongation products formed using A/H7N9 polymerase in transcription reactions. Standards (RNAs with cap1) are indicated on the left and expected products on the right. **B.(Left)** 21-mer-sd template (yellow) with T1106 single-incorporation site at position (6) and double-incorporation site at position (10-11) from the 3’ end of the native un-extended template (bracketed numbers). Expected products from transcription reactions using 12-mer capped primer (blue) and different combinations of nucleotides (black, green) and T1106 (magenta) are shown schematically. **(Right)** Gel showing elongation and stalled products formed using A/H7N9 polymerase in transcription reactions. Standards (RNAs with cap1) are indicated on the right-hand lane and expected products on the left. The stalled product upon T1106 double incorporation is visible in lane 4. **C.(Left)** 21-mer-d template (yellow) with T1106 double-incorporation site at position (10-11). Expected products from transcription reactions using 13-mer capped primer (blue) and different combinations of nucleotides (black, green) and T1106 (magenta) are shown schematically. **(Right)** Gel showing elongation and stalled products formed using FluB polymerase in transcription reactions. Standards (RNAs with cap1) are indicated on the left and expected products on the right. The stalled product upon T1106 double incorporation is visible in lane 5.

To characterise transcription reactions with T1106 we used a modified 21-mer template (denoted 21-mer-sd) that allows for one single T1106 and, later on, a double T1106 incorporation (Figure 1B left). First, we characterised the 21-mer-sd template in transcription reactions using A/H7N9 polymerase, with regular NTPs, capped 12-mer primer and 14-mer 5’ vRNA activator ‘hook’ (Figure 1B right, lane 1-3). Stalled reactions using only UTP, ATP and no CTP or GTP yielded a capped 18-mer product, the reaction lacking only GTP produced a capped 20-mer, and the reaction with all NTPs gave the full-length capped 28-mer product. Second, we utilized UTP, and T1106 instead of ATP. The transcription reaction incorporated one T1106 at the U at position (6) of the template, and then stalled, due to lack of CTP, at the same position as the reaction with UTP and ATP, resulting in the capped 18-mer product (Figure 1B right, lane 4). This is in accord with the previous studies, showing that T1106 is readily single-incorporated instead of ATP. Finally, we utilized UTP, CTP, and T1106 to replace both ATP and GTP, resulting in a transcription product of the apparent size of a capped 21-mer (Figure 1B right, lane 5), which corresponds to a stalled product upon double incorporation site of T1106 at the template (10)-UC-(11) position.

To characterise solely the effect of T1106 double incorporation, we designed a modified 21-mer-d template in which only pyrimidine bases are incorporated before the T1106 double incorporation site at (10)-UC-(11) (Figure 1C left). The 21-mer-d sequence necessitated also a modification in the corresponding 14-mer 5’ vRNA hook in order to maintain a four base-pair distal duplex in the promoter Figure 2C. Reactions were performed this time with influenza B polymerase to assess the generality of backtracking. The reaction using only UTP, CTP and stalled through lack of ATP produced a capped 19-mer, the reaction lacking only GTP produced a capped 20-mer, and the reaction with all NTPs produced the full-length capped 28-mer product (Figure 1C right, lane 1-3). The outcome of a transcription reaction utilizing UTP, CTP and T1106 is a product of an apparent size of capped 21-mer (Figure 1C right, lane 4), which correspond to a stalled product at the double incorporation site of T1106 at the template (10)-UC-(11) position.

**Figure 2.**
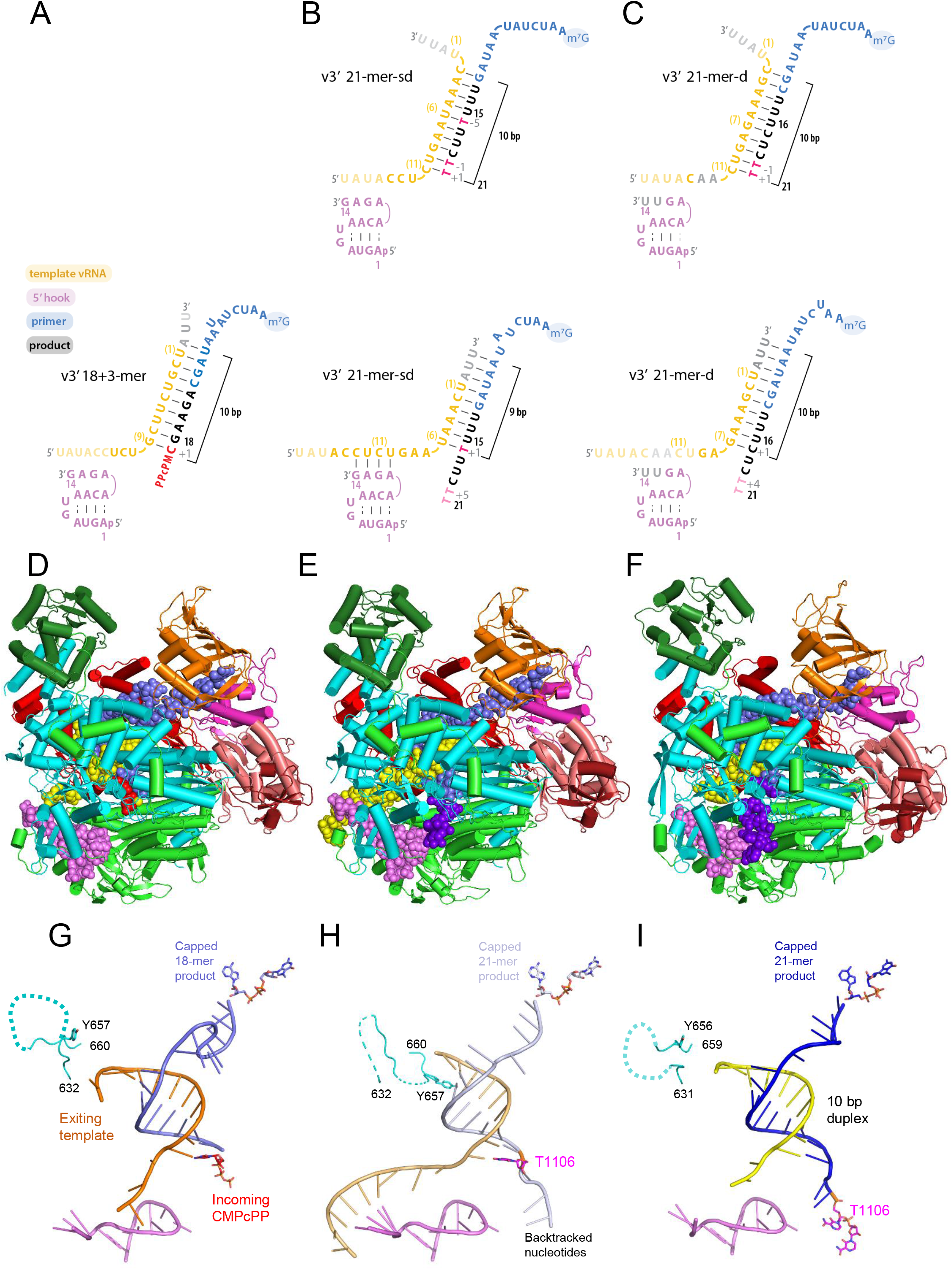
RNA configuration in the determined transcription and backtracked structures. **(A, B, C)**. Sequence and schematic secondary conformation of the RNA moieties. (**A**) A/H7N9 transcription structure stalled by CMPcPP. (**B**) T1106-stalled and backtracked A/H7N9 structure. (**C**) T1106-stalled and backtracked FluB structure. In (**B**) and (**C**), the top panel shows the presumed structure prior to backtracking. The vRNA 5′ and 3′ ends are pink and yellow respectively. The promoter duplex is disrupted except in the backtracked A/H7N9 structure where it is reformed. The bracketed numbers on the 3′ vRNA indicate distance from the native 3′ end of the template. Nucleotides added beyond the native 3′ vRNA end and the modified distal promoter are in grey. Positions with respect to the active site contain + or - signs. The capped primer is slate blue and added nucleotides black or red for analogues. Product nucleotides are numbered from the 5′ end, excluding the cap. Nucleotides not visible in the corresponding structure are in pale colours. **(D, E, F)**. Ribbon diagrams for the complete cryoEM structures. (**D**) A/H7N9 transcription structure stalled by CMPcPP. (**E**) T1106-stalled and backtracked A/H7N9 structure. (**F**) T1106-stalled and backtracked FluB structure. The polymerase colour code is PA endonuclease (forest green), PA-C (green), PB1 (cyan), PB2-N (red), PB2-midlink domain (magenta), PB2-cap-binding domain (orange), PB2-627 domain (deep pink) and PB2-NLS domain (firebrick). The RNA is coloured as in **A, B** and **C** with in **D**, the CMPcPP in red and in **E** and **F**, the observed backtracked nucleotides in purple-blue. **(G, H, I)**. Determined structures of the RNA moieties. (**G**) A/H7N9 transcription structure stalled by CMPcPP (red) with template (orange), 18-mer capped product (slate blue) and 5’ end (violet). (**H**) T1106-stalled and backtracked A/H7N9 structure with singly incorporated T1106 (magenta), template (pale orange), 21-mer capped product (pale blue) and 5’ end (violet). (**I**) the T1106-stalled and backtracked FluB structure with doubly incorporated T1106 (magenta), template (yellow), 21-mer capped product (blue) and 5’ end (violet). In each case the conformation of the priming loop residues 632-660 in A/H7N9 (631-659 in FluB) are shown in cyan, fully extruded in (**G**) and (**I**), partially extruded in (**H**) and with the position of Tyr657 (Tyr656 in FluB) highlighted.

### Structure of A/H7N9 polymerase apo-dimer and partial polymerase activation upon 5’-hook binding

Consistent with the biochemical characterisation, the cryoEM structure of full-length influenza A/H7N9 apo-polymerase revealed a C2 symmetric dimer determined at an average resolution of 2.81 Å (Table S1). Only the open, dislocated (Wandzik *et al*., 2020) dimeric core (PA/202:716, PB1/1-670, PB2/41-120) is visualised due to disorder of essentially all, flexibly linked, peripheral domains (Figure S3A). The homodimeric interface involves loops from all three subunits as previously described (Fan *et al*., 2019) and has been implicated in cRNA to vRNA replication (Chen *et al*, 2019; Fan *et al*., 2019).

The cryoEM structure of the vRNA 5’-hook bound dimer was determined at an average resolution of 3.12 Å (Table S1, Figure S3A). Of particular interest are the conformational changes that occur within the dimer upon 5’-hook binding, which have not been described before for influenza polymerase (Figure S3B-D). 5’-hook binding induces a concerted reconfiguration of PB1 peptides 23-35, 228-239 and 503-513 as well as adjacent regions of PA (e.g. side-chain of PA/W577) (Figure S3D), without changing the overall dimer structure (Figure S3A-B). This is probably initiated by distortion of PB1/23-35 to allow direct interactions of H32 and T34 with the RNA. Displacement of PB1/23-35 removes the steric hindrance to the structuring of the previously disordered fingertips loop PB1/228-239 (polymerase motif F) into its active configuration, which is locked in place, for example, by the stacking of PB1/R233 between Y24 and P28. In turn, this can only occur upon repacking of PB1/503-513. A similar allosteric activation of the fingertips loop by 5’-hook binding has been reported for La Crosse bunyavirus (Gerlach *et al*, 2015). We also observed 5’-hook bound monomers with the same structure as in the dimers (not shown). However it is only with the additional binding of the 3′ end of the promoter that the dimer is fully destabilised (Figure S1) and closure of the polymerase core and stabilisation of the peripheral domains occurs, resulting in the monomeric, RNA synthesis active form of the polymerase (Wandzik *et al*., 2020).

### Structure of an early transcription state of influenza A/H7N9 polymerase

The cryoEM structure of influenza A/H7N9 polymerase, after the incorporation of 5 nucleotides and stalled with the non-hydrolysable analogue CMPcPP at the +1 position (Figure 2A), was determined at an average resolution of 2.48 Å (Table 1). The structure (Figure 2D), which has very good to reasonable density for the entire heterotrimer (Figure S2A), depicts an early elongation state after the transition from initiation is complete. It is remarkably similar to the equivalent elongation structure previously determined for bat influenza A/H17N10 (PDB:6T0V, (Wandzik *et al*., 2020), RMSD 0.72 Å for PB1/1-560), as well as to the less homologous influenza B polymerase elongation structure (PDB:6QCT, (Kouba *et al*., 2019) RMSD 0.84 Å PB1/1-560). Like these structures, the polymerase has slightly opened to accommodate the full 9-mer template-product duplex in the active site cavity (positions -1 to -9) and the priming loop is fully extruded (Figure 2G), allowing the template to enter the exit channel after strand separation enforced by PB2/Tyr205 (from the helical lid) and PB1/R706. The most significant differences to the bat influenza A early elongation structure are, firstly, the incoming nucleotide, here CMPcPP compared to UMPnPP in PDB:6T0V, has an alternative triphosphate conformation that does not support the catalytic configuration of Mg(A) and Mg(B) (Figure 3A); instead there is a well-defined Mg(A’) in the inactive position close to PB1/E491 (Wandzik *et al*., 2020). Secondly, the distal end of the PB1/β-ribbon is poorly ordered and there is no clear formation of a three-stranded β-sheet together with residues PB1/667-681 (Kouba *et al*., 2019; Wandzik *et al*., 2020). Thirdly, unlike the bat influenza A and influenza B elongation structures, the entire 18-mer capped primer/product is visible. To accommodate nucleotides 1-9 of the capped product between the cap and the product-template duplex, the RNA takes up a very particular conformation (Figure S4A). Cap-proximal bases A2, U3 and C4 are stacked on each other, with PB2/Arg144 (aliphatic part stacks on C4), Arg213 (fortuitous base-specific interactions with C4) and Arg436 (packs against U3 ribose and hydrogen bond to the 2’ OH of A2) playing key roles in stabilising this configuration. This is followed by a tight turn (with C4 ribose interacting with A7 phosphate) and then a distinct stacking of bases of U5, A6, A8 and A9, with U5 packed on PB2/Arg423 and U7 being bulged out (Figure S4A). Interestingly, the cap-proximal nucleotides 2-4 are orientated quite differently in the A/H7N9 structure compared to both bat A/H17N10 (PDB:6T0V elongation, PDB:6EVJ pre-initiation) and FluB (PDB:6QCT elongation, PDB:6QCX pre-initiation) capped RNA bound structures, which have similar configurations (Kouba *et al*., 2019; Pflug *et al*, 2018; Wandzik *et al*., 2020)(Figure S4B). This is likely attributable to the substitution in human/avian influenza A of PB2/H432 by Y432 and Y434 in respectively bat A/H17N10 and FluB polymerases. Capped primer nucleotides 2-4 are stacked on each other in all these cases, but in bat A/H17N10 and FluB the tyrosine defines the orientation by stacking on the base of nucleotide 2, with PB2/Arg217 stacking on the fourth base in the case of FluB (Figure S4B). Comparing more specifically A/H7N9 and bat FluA, where essentially all other residues in the vicinity are conserved, the plane of the side-chain of H432 in A/H7N9 is perpendicular to that of Y432 in bat A/H17N10, allowing it to hydrogen bond to the first phosphate of the cap-triphosphate as well as to the 2’ OH of the second nucleotide. Furthermore, H432 does not stack with nucleotides 2-4, which appear to be positioned by the arginine residues mentioned above. It cannot be ruled out that other factors influence the particular conformation of the capped primer/product, for instance the sequence (here m^7^GTP-AAUC for A/H7N9 and m^7^GTP-GAAU for bat A/H17N10 and FluB). However, in the A/H5N1 pre-initiation structure with bound promoter and capped RNA, the sequence of the primer (m^7^GTP-AAUC) and the position of H432 is the same as in A/H7N9, but nucleotides 2-4 are arranged quite differently and not stacked on each other. Furthermore, in the backtracked structures (see below), for the case of A/H7N9, the stacking of the cap-proximal nucleotides 2-4 is preserved, but in the FluB structure only the second primer base stacks on the tyrosine and bases 3-4 are in a different orientation and sandwiched between PB2/R217 and R438 (Figure S4B). This suggests that the length of the capped primer/product and the trajectory required to connect it to the active site in initiation, or to the template-product duplex in elongation, determines how compact or, alternatively, stretched, the capped oligomer conformation is, with PB2 residues, in particular arginines (e.g. for A/H7N9, PB2/R144, R213, R423, R436), being able to accommodate to a wide variety of alternatives by stacking and/or hydrogen bonding.

**Table 1.** CryoEM structure determination and validation statistics of elongation and backtracked structures.

| Name of structure | A/H7N9-Elongation | A/H7N9-Backtracked-T/U+1 | B/Mem-Backtracked |
| --- | --- | --- | --- |
| PDB ID | **7QTL** | **7R0E** | **7R1F** |
| EMDB ID | **EMD-14144** | **EMD-14222** | **EMD-14240** |
| **Data collection and processing** | EMBL DATA1 | ESRF DATA 1 | EMBL DATA2 |
| Microscope | FEI Titan Krios | FEI Titan Krios | FEI Titan Krios |
| Voltage (kV) | 300 | 300 | 300 |
| Camera | Gatan K2 Quantum | Gatan K2 Quantum | Gatan K3 Quantum |
| Magnification | 165,000 | 165,000 | 105,000 |
| Nominal defocus range (μm) | 0.5 – 1.8 | 0.4 – 2.1 | 0.5 – 2.5 |
| Exposure time (s) | 5 | 7.2 | 1.3 |
| Electron exposure (e-/Å<sup>2</sup>) | 33.4 | 48 | 46.2 |
| Number of frames collected (no.) | 40 | 51 | 40 |
| Number of frames processed (no.) | 22 | 24 | 20 |
| Pixel size (Å) | 0.8127 | 0.827 | 0.822 |
| Micrographs (no.) | 3214 | 7336 | 13747 |
| Total particle images (no.) | 1,238,425 | 1,472,891 | 8,451,594 |
| **Refinement** |  |  |  |
| Particles per class (no.) | 116,539 | 98,908 | 328,180 |
| Map resolution (Å), 0.143 FSC | 2.48 | 2.51 | 2.58 |
| Model resolution (Å), 0.5 FSC |  |  |  |
| Map sharpening *B* factor (Å<sup>2</sup>) | -20.84 | -67.66 | -77.0 |
| Map versus model cross-correlation (CCmask) | 0.83 | 0.87 | 0.84 |
| **Model composition** |  |  |  |
| Non-hydrogen atoms | 18397 | 18554 | 18387 |
| Protein residues | 2152 | 2162 | 2177 |
| Nucleotide residues | 44 | 49 | 42 |
| Water | 120 | 114 | 6 |
| Ligands | 4 | 4 | 6 |
| ***B* factors (Å<sup>2</sup>)** |  |  |  |
| Protein | 36.77 | 61.83 | 64.46 |
| Nucleotide | 37.92 | 59.74 | 59.64 |
| Ligand | 29.45 | 58.35 | 146.15 |
| Water | 22.33 | 46.23 | 35.55 |
| **R.M.S. deviations** |  |  |  |
| Bond lengths (Å) | 0.003 | 0.003 | 0.003 |
| Bond angles (°) | 0.611 | 0.474 | 0.466 |
| **Validation** |  |  |  |
| MolProbity score | 2.01 | 1.33 | 1.74 |
| All-atom clashscore | 8.57 | 5.25 | 6.73 |
| Poor rotamers (%) | 3.98 | 1.15 | 1.89 |
| **Ramachandran plot** |  |  |  |
| Favored (%) | 97.56 | 98.04 | 97.09 |
| Allowed (%) | 2.39 | 1.96 | 2.86 |
| Outliers (%) | 0.05 | 0 | 0.05 |

**Figure 3.**
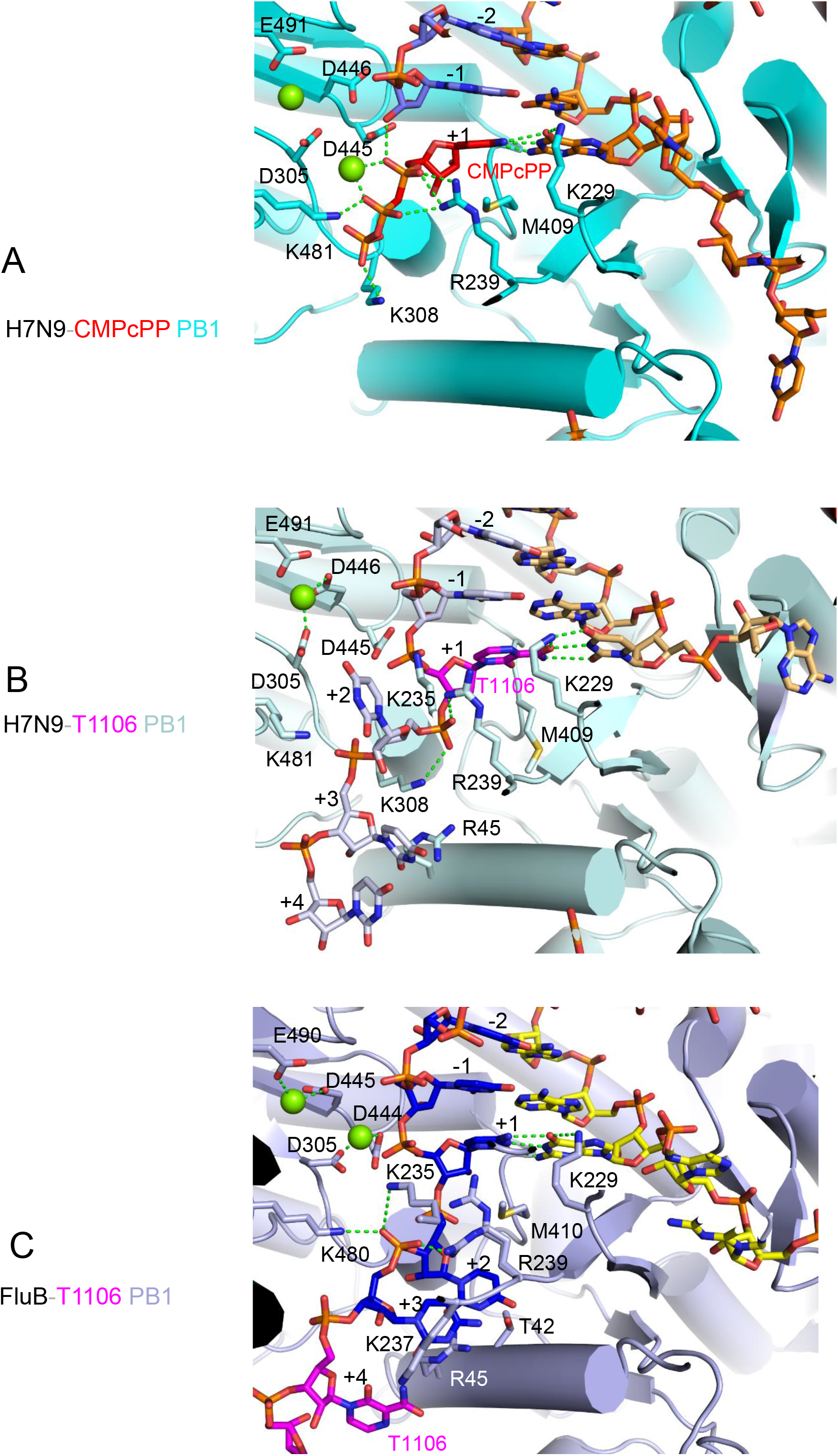
Active site details for elongation and backtracked structures. **(A, B, C)**. Key features and interactions within the polymerase active site region of each determined structure. (**A**) A/H7N9 transcription structure stalled by CMPcPP (red) with template (orange), product (slate blue) and PB1 cartoon and residues (cyan). (**B**) T1106-stalled and backtracked A/H7N9 structure with singly incorporated T1106 (magenta) at the +1 position, template (pale orange), product (pale blue) and PB1 cartoon and residues (pale cyan). **(C)** T1106-stalled and backtracked FluB structure with template (yellow), product (blue) and PB1 cartoon and residues (light blue). Indicated nucleotide positions are relative to +1 at the active site. Magnesium ions are green spheres and putative hydrogen bonds dotted green lines.

### Structure of stalled and backtracked state of influenza A/H7N9 polymerase

The cryoEM structure of influenza A/H7N9 polymerase, stalled and backtracked by five nucleotides after double incorporation of T1106 and with singly incorporated T1106 at the +1 position was determined at an average resolution of 2.51 Å (Table 1, Figure 2B, 2E). The high resolution allows unambiguous assignment of the RNA sequence (Figure S2B). According to the biochemistry, single incorporation of T1106 at U6 of the template followed by double incorporation at template positions (10)-UC-(11) leads to a stalled product corresponding to a capped 21-mer. This product corresponds to an addition of nine nucleotides to the 12-mer capped primer, with T1106 incorporated at positions 16 and 20-21. If there was no backtracking (Figure 2B top), one might expect a stalled elongation state of the polymerase similar to that described above. This would have a 10-mer template-product duplex in the active site cavity with T1106 at the -5, -1 and +1 positions of the product, the priming loop fully extruded, the leading four nucleotides of the template in the exit channel and the promoter duplex disrupted. However, this state appears to be unstable, as it is not observed on the cryoEM grid. Instead, backtracking by five nucleotides of the entire RNA system is observed presumably stopping when a stable state is reached. This backwards translocation results in a 9-mer template-product duplex in the active site cavity with T1106 at the +1, +5 and +6 positions of the product, no template nucleotides in the exit channel, partial reinternalization of the priming loop and reformation of the promoter duplex, but with an unusually short connection to the +1 position (Figure 2B, 2H). The backtracked 21-mer capped RNA product is partitioned between only seven nucleotides connecting to the m^7^G cap bound in the cap-binding site, forming a direct connection (i.e. eliminating the bulge seen in the elongation structure, Figure S4C) to nine nucleotides in the template-product duplex and five nucleotides in the +2 to +6 positions projecting back into and filling the incoming NTP channel. Of the backtracked product, only the 17-UU-18 in the +2 and +3 positions are clearly defined by the cryoEM density. The base of U17 is sandwiched between PB1/K253 (motif F) and PB1/K481 (motif D) and its phosphate interacts with PB1/K308 (motif A) and PB1/R239 (motif F) (Figure 3B). U18 base contacts PB1/K237 (motif F) and PB1/R45 and its phosphate is close to the putative position of the gamma phosphate of an incoming NTP (Figure 3B, 4B). Lower resolution density encompasses C19 at the +4 position, but not beyond, presumably because of disorder of the extruded and untethered 3’ end of the product. Thus, there is unfortunately no direct structural conformation for the double incorporation of T1106, although the biochemistry shows that this is the case.

Compared to the A/H7N9 elongation structure, there is an axial shift (base displacement) of about 1 Å downwards of the template strand in the product-template duplex, and a corresponding to 1.7-2.3 Å helical change in phosphate positions (Figure 4A). This necessitates a correlated change in backbone of the PB1/125-128 to accommodate the shifted phosphate of template nucleotide (A5) in the -1 position. In the normal elongation structures, PB1/Met409 (motif B) stacks under the base at the product +1 (Kouba *et al*., 2019). In the backtracked structure, motif B backbone is slightly altered to allow reorientation of the Met409 sidechain so that it does not clash with the product base in the +1 position (T1106) (Figure 4A).

**Figure 4.**
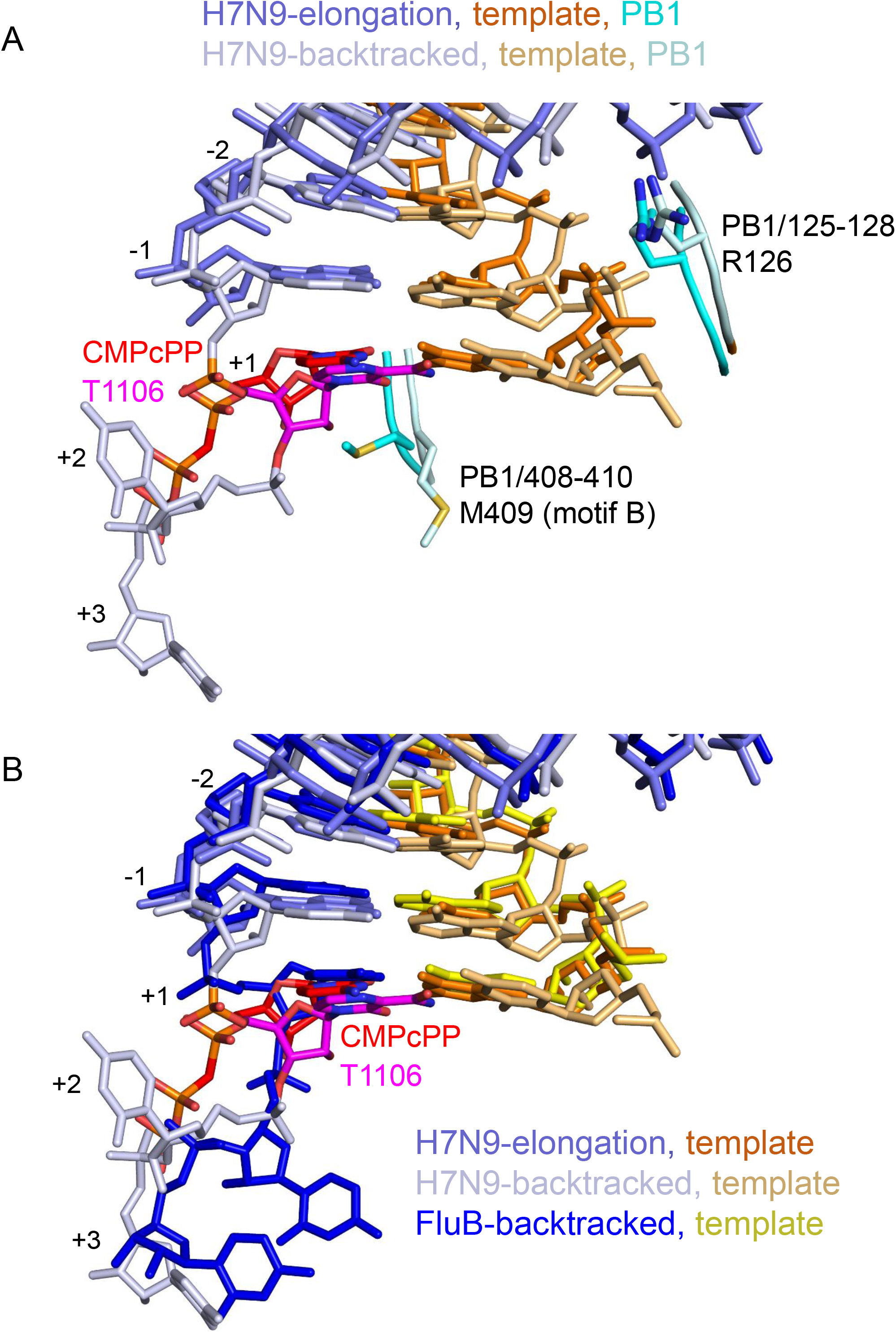
Comparison of the RNA configuration between elongation and backtracked structures. **A)** Comparison of the position of the RNA in the active site region of the A/H7N9 transcription structure (template orange, product slate-blue, PB1 cyan), stalled with CMPcPP (red) and backtracked structure (template pale-orange, product light-blue, PB1 light cyan) stalled with T1106 (magenta) after superposition of PB1 subunits (RMSD 0.45 Å for superposition of PB1/1-560). To accommodate the small downward shift of the duplex RNA in the backtracked structure requires local adjustment of the rotamer of PB1/M504 (motif B) and the position of PB1/R126. **B)** Comparison of the position of the RNA in the active site region of the A/H7N9 backtracked structure (template pale-orange, product light-blue, PB1 light cyan, T1106 magenta) with that in the FluB backtracked structure (template yellow, product blue) after superposition of the PB1 subunits (RMSD 0.81 Å for superposition of PB1/1-560). The figure highlights the shifted axial position of the duplex and the different disposition of backtracked nucleotides in the +2 and +3 positons.

Two other consequences of the backtracking are reformation of the promoter duplex and partial reinternalization of the priming loop. Backtracking by five nucleotides allows the reformation of the same 4 base-pair promoter duplex between 5’ end nucleotides 11-14 and 3’ template nucleotides (10)-(13), as expected for the pre-initiation state. However, in the pre-initiation state the template takes an indirect path to the active site -1 position requiring six nucleotides, whereas in the backtracked structure there is a taut connection requiring only three nucleotides (Figure S4D). The backtracked conformation therefore mimics a late stage in the initiation to elongation transition, in which the template has translocated by three nucleotides through the active site, shortening and straightening the connection to the duplex region of the promoter, but not breaking the duplex. Such a state has been observed in recent structures of La Crosse bunyavirus polymerase (Arragain *et al*, 2022). Consistent with this, the polymerase has opened to accommodate the product-template duplex in the active site cavity but the priming loop is only partially extruded. The only other structure with a partially extruded priming loop was for influenza B polymerase at an earlier stage in the transition. This had five base-pairs in the active site cavity and most of the priming loop was still internalised with only residues 636-640 disordered in the solvent (Kouba *et al*., 2019). Here, there are nine base-pairs in the product-template duplex and the priming loop is in a different conformation where it is pushed further out but not fully (Figure 2H). In particular, priming loop residue PB1/Y657 is intercalated between PB1/V525 and the ribose of template nucleotide U(−2) (with its OH group hydrogen bonding to 2’OH of A(−1), forcing displacement of the distal part of the product-template duplex away from its position when the priming loop is fully extruded (Figure S4C). Finally, the active site is in a non-catalytic configuration with a single hydrated magnesium co-ordinated by PB1/D305, D446 and E491 (Figure 3B).

It is generally assumed that T705 will form canonical (i.e. Watson-Crick like) base-pairs with cytidine and uridine (Figure 5A) (Jin *et al*., 2013; Wang *et al*., 2021). However there are two possible base-pairs that T705 or T1106 can form with uridine, a canonical base-pair resembling a Watson-Crick A:U and an alternative wobble base-pair resembling G:U (Figure 5B). At the +1 position, U(6) of the template forms a wobble base-pair with the singly incorporated T1106 at nucleotide 16 of the product (Figures 2B, 2H, 3B, 5). Whereas in a Watson-Crick A:U base-pair, the two hydrogen bonds are to the N3 and O4 of uridine, here we observe that the rotatable amide of T1106 hydrogen bonds the with O2 and N3 of U(6), similar to a G:U wobble base-pair (Figure 5B). PB1/K229 interacts with the O4 of U(6) and more distantly (3.2 Å) with the oxygen of the T1106 amide (Figure 5B). There is also a water model in the same plane the other side of the base-pair, interacting with the O2 of U(6), the amide NH2 of T1106 as well as the carbonyl oxygen of PB1/M409 and backbone of A242 (Figure 5B). The K229R substitution has been identified as imparting partial resistance to T705, whilst at the same time reducing polymerase activity (Goldhill *et al*., 2018). The nature of the T1106:U base-pair is discussed further below.

**Figure 5.**
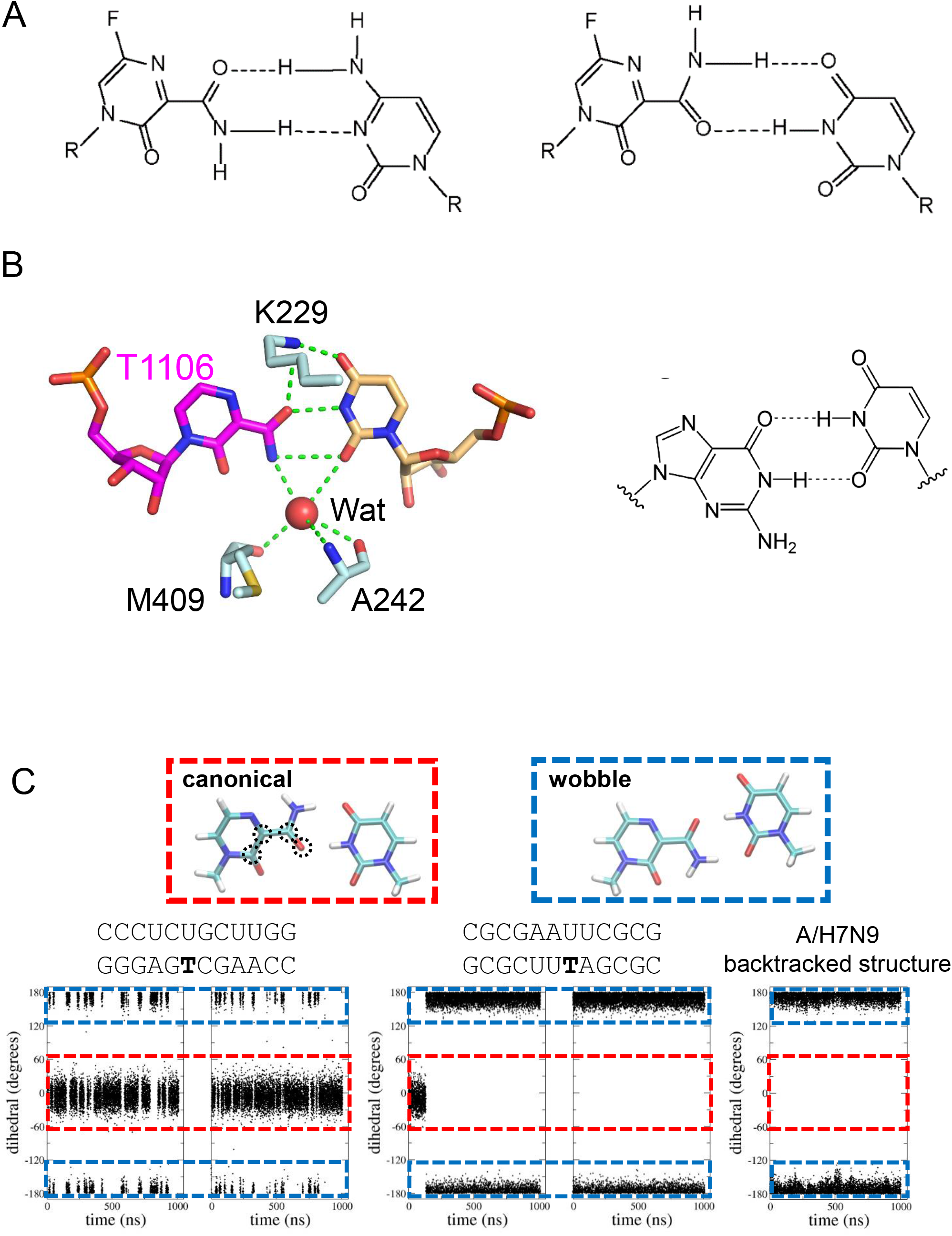
Canonical and wobble T1106:U base-pairing. **A)** Canonical T705:C (left) and T705:U (right) base-pairs from (Jin *et al*., 2013). **B)** Left: Observed T1106:U wobble base-pair at the +1 position of the A/H7N9 backtracked structure showing interactions with PB1/K229 and a water molecule. Right: Wobble G:U base-pair. **C**). Evolution of the T1106 amide group dihedral angle (defined by the circled atoms in the molecular model of the canonical base-pairing) during the MD simulations of RNA duplexes (left and center) and of the backtracked A/H7N9 structure (right). For each RNA duplex the sequence is shown and simulations starting from either canonical or wobble U:T1106 base pairing were performed (left, right respectively).

### Structure of stalled and backtracked state of influenza B/Memphis polymerase

The cryoEM structure of influenza B polymerase, stalled and backtracked by four nucleotides after double incorporation of T1106 was determined at an average resolution of 2.58 Å (Table 1, Figure 2C,2F). The high resolution allows unambiguous assignment of the RNA sequence (Figure S2C). According to the biochemistry, double incorporation at template positions (10)-UC-(11) leads to a capped 21-mer product, corresponding to the addition of 8 nucleotides to the 13-mer capped primer, with T1106 incorporated at positions 20-21. Without backtracking, one would expect a stalled elongation state of the polymerase that would have a 10-mer template-product duplex in the active site cavity with T1106 at the -1 and +1 positions of the product, the priming loop fully extruded, the leading four nucleotides of the template in the exit channel and the promoter duplex disrupted (Figure 2C top). Instead, backtracking by four nucleotides of the entire RNA system is observed. This results in a 10-mer template-product duplex in the active site cavity (abutted at the distal end against PB2/Y207 of the helical lid), with T1106 at the +4 and +5 positions at the 3’ end of the product and no template nucleotides in the exit channel (Figure 2C bottom, 2F, 2I). However, unlike the backtracked A/H7N9 structure, the priming loop remains fully extruded and the promoter duplex does not reform (Figure 2I). The backtracked 21-mer capped RNA product is partitioned between seven nucleotides connecting to the m^7^G cap bound in the cap-binding site (see Figure S4B for the conformation of the cap-proximal nucleotides), ten nucleotides in the template-product duplex and four nucleotides in the +2 to +5 positions projecting back into the incoming NTP channel (Figure 2C, 2I). Of the backtracked product, 18-UC-19 in the +2 and +3 positions are clearly defined by the cryoEM density, but they are in quite different positions than seen in the A/H7N9 backtracked structure (Figure 3C). The base of U18 is stacked on PB1/T42 and R45. The base of C19 is partially stacked on that of U18 and PB1/K237 on the other side; its phosphate interacts with PB1/K235, R239 and K480. There is clear density in blurred maps for the T1106 at position 20 and the backbone of T1106 at position 21, but not good enough to confirm the base identities. The observation of backtracking upon consecutive T1106 incorporation also in the case of FluB polymerase, where there is no prior single incorporation, strongly suggests that the backtracking in the case of A/H7N9 polymerase is due to the double incorporation and not related to the prior single incorporation event.

The axial position of the product-template duplex in the FluB backtracked structure is closer to that of the transcription elongation structure (Figure 4B). Although the FluB density suggests alternate conformations of the motif B loop, the main conformation maintains PB1/M410 (equivalent to M409 in A/H7N9) stacked under the product base in the +1 position (Figure 3C), whereas in the A/H7N9 backtracked structure, the lower positon of the base forces the M409 into a different, non-clashing rotamer (Figure 4A).

## Discussion

T705 is thought to have a dual mode of action, lethal mutagenesis in the case of single incorporation and chain termination in the case of consecutive incorporation (Jin *et al*., 2013; Wang *et al*., 2021). Similarly, it is shown here, biochemically and structurally, that T1106 can be readily incorporated singly without termination but induces chain termination, without further addition, after two successive incorporations. Possible explanations for chain termination are that after double incorporation (a) the RNA does not translocate in the forward direction, thus preventing NTP entry at the +1 position, (b) the RNA translocates but the next incoming NTP cannot bind or (c) the RNA translocates, the next NTP binds but does not react. A previous study of this phenomenon using biochemical assays and molecular dynamics (MD) simulations concluded that consecutive incorporation of T705 disrupts the base stacking of the product strand and destabilises the active site, such that the next incoming NTP is bound but not productively and thus not incorporated, corresponding to explanation (c) (Wang *et al*., 2021). However the MD results were based on models in which it was already assumed that translocation in the forward direction had occurred. Here we provide new insight into the structural basis for chain termination with the observation that in two distinct cases, involving FluA and FluB polymerases, double incorporation of T1106 leads to RNA backtracking, which can be considered as an extension of explanation (a). Furthermore, we observe an instance of a singly incorporated T1106:U base-pair at the +1 position of the A/H7N9 backtracked structure. The high resolution of our structure enables unambiguous assignment of a T1106:U wobble base-pair, rather than canonical A:U like base pair (Figure 5). Interestingly, recent quantum mechanical calculations of the relative stability of all possible base-pairs involving T705 show that the T705:U wobble and canonical base-pairs have binding energies of respectively -11.45 and -9.15 kcal/mol showing that the wobble base-pair is 2.3 kcal/mol more stable (Fig S1 of (Jena, 2020)). We have made similar calculations and obtain an extra stability of the wobble base pair of 2.70 and 2.09 kcal/mol for respectively T705 and T1106 (Figure S5A). Further, we have assessed by microsecond long equilibrium molecular dynamics the nature of the T1106:U base-pair over time when embedded in an RNA double-helix. With two different flanking sequences, we find different behaviour including long-term stability of the wobble base pair or rapid flipping between the two types (Figure 5C). This suggests that due to the relative small energy differences, the local environment can play a determining role for the configuration of the T1106:U base-pair. Indeed, the wobble base pair was stably maintained in the A/H7N9-backtracked-T/U+1 complex throughout additional microsecond-long MD simulations (RMSD value for the heavy atoms of T1106:U base pair of 1.0 ± 0.2 Å, Figure S5B) This is likely due to the steady interaction of the drug with K229 (Figure S5C). In the light of these considerations, our observation of a T1106:U wobble base-pair at the +1 position in the back-tracked structure is perfectly plausible. However, in other situations, for instance an incoming T1106-TP opposite a U in the template at the +1 position, it could be different. To gain further insight into this, we examined other published polymerase structures that contain T705, there being none containing T1106. Those containing T705 are all recent studies of SARS-CoV-2 polymerase. There are two pre-incorporation structures of SARS-CoV-2 polymerase with incoming T705-TP opposite a C at the +1 position in the template (PDB:7AAP (Naydenova *et al*., 2021), PDB:7CTT (Peng *et al*., 2021)). In both cases, the canonical T705:C base-pair is observed, consistent with this being the only energetically stable form (c.f. Fig S1 of (Jena, 2020)). There is also a deposited structure (PDB:7DFG), but without publication, in which a post-incorporation, pre-translocation T705:U base-pair is modelled in canonical form, together with pyrophosphate. However the cryoEM density (EMD-30663) for the T705 moiety is very weak, indicating partial occupancy and furthermore, there appear to be errors in assignment of some other bases in the template-product duplex, making the register of the stalled reaction unclear.

In the MD simulations of T705 double incorporation by influenza polymerase (Wang *et al*., 2021) it was assumed, without discussion, that the incoming T705:U base-pair would be canonical. Nevertheless, this MD study strongly suggests that two successive incorporations of T705, but not just one insertion, destabilises the base stacking of the product strand as well as hindering productive binding of the next NTP. Whether this is partly due to the possibility of flipping between canonical and wobble T705:U base pairs is not clear. However, our structural results suggest that what happens next is backtracking, with the primary driving force being to remodel the overall RNA into a more stable configuration, as also evidenced by our MD simulations (RMSD value for the heavy atoms of RNA-RdRp complex of 2.8 ± 0.2 Å, Figure S5D). Backtracking achieves this by extruding the consecutive T705/T1106 bases into a single stranded region and maximising the number of stable Watson-Crick base pairs in duplex regions. However, the exact post-backtracking configuration likely depends on a complicated energetic balance that results in an idiosyncratic response. In the two examples we present, A/H7N9 and FluB, the presumed structure prior to backtracking (Figure 2BC, top) is very similar as stalling occurs at the same point in the template with the capped product in both cases being a 21-mer. The most significant difference is the occurrence of the single T1106:U base pair in the middle of the product-template duplex in the A/H7N9 case compared to a U:A in the FluB case. However, backtracking leads to two quite different structural outcomes. For FluB, backtracking by only four nucleotides leaves the maximal 10 base-pair duplex (stabilised at the distal end by packing against the helical lid) in the active site cavity and the priming-loop remains extruded. For A/H7N9, a stable configuration is reached after backtracking by five nucleotides leaving a 9 base-pair duplex in the active site cavity with a wobble T1106:U base pair at the +1 position. The distal end of the slightly shorter product-template duplex is stabilised by the partially reinserted priming loop, and loss of the tenth duplex base pair is further compensated by reformation of the four base pair promoter. Other factors such as strength of protein-RNA interactions and conformation of the product RNA connecting to the cap-binding site (Figure S4) may also play a role in defining the final backtracked conformation.

Most *in vitro* studies of T705 inhibition, like those on T1106 described here, have detected chain termination in very early elongation. This is physiologically relevant because the 3’ end of the vRNA template is pyrimidine-rich with several opportunities for double incorporation at a very early stage if the concentration of the nucleotide analogue is high enough (Wang *et al*., 2021). On the other hand, early elongation is particular in that it coincides with the initiation to elongation transition, which involves promoter duplex breaking, extrusion of the priming loop, opening of the polymerase and first establishment of the 10 base pair product-template duplex (Kouba *et al*., 2019). In this context, chain termination and backtracking can lead to idiosyncratic conformations including partial reversal of the initiation to elongation transition as described above. To characterise the effect of T1106 double incorporation during steady-state elongation, we designed a 56-mer mini vRNA (i.e. with connected 3’ and 5’ ends) with a purine-rich template that contains a double incorporation site for T1106 at the template position (30)-CC-(31) (Figure S6). Due to the loop design and lack of poly-adenylation signal, the canonical transcription stop site is expected to be at position A40 (i.e. A17 from the template 5’-end)(Wandzik *et al*., 2020). We then performed different RNA synthesis reactions giving sufficiently long products that the 3’ end of the translocating template will have docked into the secondary 3’ end binding site (Wandzik *et al*., 2020). First, we characterised this 56-mer template in transcription reactions primed with an un-capped 6-mer primer and regular NTPs (Figure S6, lanes 1,4 and 5). The reaction using only UTP, CTP and stalled through lack of GTP produced a 30-mer product, addition of UTP, CTP and GTP as well as all NTPs produced the expected full-length 40-mer transcript. Next, we utilized UTP and CTP in combination with 2’FdGTP (2’-deoxy-2’-fluoroguanosine triphosphate), which was previously shown to be a non-obligate chain terminator for influenza virus polymerase (Tisdale *et al*, 1995). Indeed, upon 2FdGTP incorporation, the transcription progressed by one base in comparison to UTP/CTP product, and produced 31-mer product (Figure S6, lane 2). Finally, the reaction containing UTP, CTP and T1106, produced a 32-mer product, which correspond to a stalled product at the double incorporation site of T1106 at the template CC position (30-31) (Figure S6, lane 3). Unfortunately, examination of cryoEM grids made with the latter reaction did not yield any classes corresponding to stalled polymerase, only the pre-initiation state. Thus, we were unable to determine whether backtracking took place upon T1106-induced stalling in the steady-state elongation situation before product dissociation.

T1106-induced stalling and backtracking of poliovirus polymerase has recently been deduced from single-molecule experiments using magnetic tweezers (Dulin *et al*., 2017)} and these observations have recently been extended to enterovirus A-71 polymerase (Janissen *et al*., 2021) and SARS-CoV-2 (Seifert et al., https://www.biorxiv.org/content/10.1101/2020.08.06.240325v2). For poliovirus it was shown that incorporation of T1106 leads to prolonged pausing for tens to thousands of seconds often associated with back-tracking by tens of nucleotides, although the system can eventually recover and elongation proceed again. It is intriguing that backtracking has now been observed in several quite different viral polymerases, uniquely for T1106 and not for other nucleoside analogues, such classical chain terminators (e.g. 3’-dATP) or other mutagens (e.g. ribavirin). It will be interesting to see whether the pyrimidine mimic and mutagen, NHC-TP (β-D-N4-hydroxycytidine-TP), which has been developed as an anti-influenza compound (Toots *et al*, 2019) and as the anti-SARS-CoV-2 (Sheahan *et al*, 2020) drug Molnupiravir, also induces backtracking.

Backtracking in response to misincorporation of normal NTPs or nucleoside analogues is used by cellular (Cheung & Cramer, 2011), bacterial (Abdelkareem *et al*., 2019) and coronavirus (Malone *et al*., 2021; Robson *et al*, 2020) RNA polymerases as a mechanism of error correction by proof reading. The polymerase active site can theoretically use pyrophosphate to catalyse the hydrolysis of the backtracked product RNA, thus creating a new product 3’ end that can be elongated, but this reaction is slow under physiological conditions. Specific excision mechanisms and factors have therefore evolved to remove the backtracked nucleotides e.g. TFIIS in eukaryotes (Cheung & Cramer, 2011), GreA and GreB in bacteria (Abdelkareem *et al*., 2019), or the coronavirus NSP14 exonuclease (Liu *et al*, 2021). For influenza polymerase there is no evidence to date that such a mechanism exists and our observed backtracked structures are sterically incompatible with pyrophosphate-mediated hydrolysis of the RNA at the polymerase active site.

## Supporting information

Supplemental Table S1, Supplemental Figures and Legends, Supplemental CryoEM Methods

## Acknowledgments

We thank the EMBL Grenoble Eukaryotic Expression Facility (Alice Aubert) and the EMBL-ESRF-ILL-IBS Partnership for Structural Biology (PSB) biophysical platform (Caroline Mas). We thank Felix Weis and Wim Hagen for access to the Titan Krios at EMBL Heidelberg, Wojtek Galej and Erika Pellegrini for access to the Glacios at EMBL Grenoble, and Aymeric Peuch and Thomas Hofmann for help using the joint EMBL-IBS or EMBL Heidelberg computer clusters, respectively. We acknowledge the European Synchrotron Radiation Facility (ESRF) and PSB for access to the Titan Krios CM01 and thank Michael Hons and Eaazhisai Kandiah for assistance with data collection.

## Grant support

This work was supported by ANR grant (ANR-18-CE11-0028) to SC. This work used the platforms of the Grenoble Instruct-ERIC Center (ISBG ; UMS 3518 CNRS-CEA-UGA-EMBL) within the Grenoble Partnership for Structural Biology (PSB), supported by FRISBI (ANR-10-INBS-05-02) and GRAL, financed by the Université Grenoble Alpes graduate school (Ecoles Universitaires de Recherche) CBH-EUR-GS (ANR-17-EURE-0003).

## Author contributions

V.S. and P.D. expressed and V.S., P.D. and T.K. purified A/H7N9 and B/Memphis polymerases. J.H and C.M synthesized and provided T1106-TP. A.D. and T.K performed transcription assays. A.D. and T.K prepared cryoEM grids, A.D. and T.K. collected cryoEM data, T.K. performed cryoEM image processing and 3D reconstruction. S.C. built and refined atomic models. E D. and P.V. performed and analyzed the molecular dynamics computations, supervised by M.D.V. T.K and S.C conceived and supervised the project and prepared the manuscript and figures with input from all other authors.

## Declaration of interests

The authors declare no competing interests.

## MATERIALS and CORRESPONDANCE

Further information and requests for resources and reagents should be directed to and will be fulfilled by the Lead Contact, Stephen Cusack.

## METHODS

### Expression and purification

Influenza A/Zhejiang/DTID-ZJU01/2013 (H7N9) polymerase heterotrimer was expressed from a codon-optimized synthetic construct containing His8x-PA (Uniprot: M9TI86) – PB1 (Uniprot:M9TLW3) – PB2-StrepII (Uniprot: X5F427) cloned into pET-DUAL vector for co-expression under the control of the polH (for PA and PB1) and p10 (for PB2) promoters. The A/H7N9 polymerase was produced using the baculovirus expression system in HighFive insect cells, which were collected after 48-60h post-infection by centrifugation, re-suspended in buffer A (50 mM HEPES/NaOH pH8, 500 mM NaCl, 10% (v/v) glycerol) supplemented with protease inhibitors (Roche, complete mini, EDTA-free, leupeptin, pepstatinA), and lysed by sonication on ice. The cell extract was then cleared by centrifugation (30 min, 4 °C, 35,000*g*) and ammonium sulphate added to the supernatant with ratio 0.6 g/ml lysate for protein selective precipitation and recovery. The recombinant protein was then collected by centrifugation (30 min, 4 °C, 70,000 *g*) and re-suspended in buffer A and the procedure repeated twice. H7N9 polymerase was then purified from the soluble fraction via tandem affinity chromatography: Ni-sepharose metal binding, followed by strep-tactin (IBA, Superflow), using buffer A (supplemented with 400 mM imidazole for Ni affinity) as mobile phase in both cases. Protein-containing fractions were pooled and diluted with an equal volume of buffer B (50 mM HEPES/NaOH pH 7.5, 10% (v/v) glycerol) before loading on to a third affinity column (HiTrap Heparin HP, GE Healthcare). A step gradient (25-50-75-100%) of buffer C (buffer B supplemented with 1 M NaCl) was applied, and polymerase was eluted as single species at 600 mM NaCl. Finally, the protein was concentrated with Amicon® Ultra-15 (50 KDa cutoff) to ~6 μM, flash-frozen and stored at -80C.

The influenza B/Memphis/13/03 (FluB) polymerase self-cleaving polyprotein heterotrimer construct was expressed in High Five insect cells as described previously (Reich *et al*, 2014). Frozen cell pellets were re-suspended in lysis buffer (50 mM Tris-HCl, 500 mM NaCl, 10% glycerol, pH 8) containing protease inhibitors (Roche, complete mini, EDTA-free). Following lysis by sonication and centrifugation at 20,000 r.p.m. (JA20/Beckman Coulter) for 45 min at 10°C, the supernatant was precipitated by ammonium sulphate (0.5 g ml^−1^) and centrifuged at 45,000 r.p.m. for 45 min at 10°C (45Ti/Beckman Coulter). The pellet was re-dissolved in lysis buffer, and finally re-centrifuged at the same settings. Cleared supernatant was incubated with nickel resin (His60 NiNTA, Clontech) for one hour at 10° C. Protein was eluted with lysis buffer supplemented with 500 mM imidazole and loaded on a Strep-Tactin matrix (Superflow, IBA). Elution was performed with 2.5 mM d-desthiobiotin in low salt buffer (50 mM Tris pH 8, 250 mM NaCl, 10% (v/v) glycerol). Pooled FluB polymerase fractions were filtered with 0.22 μm filter and loaded on a heparin column (HiTrap Heparin HP, GE Healthcare). Elution was performed with a gradient using buffers A and B (2 mM TCEP, 50 mM HEPES, pH 7.5, 150 mM (A) or 1 M NaCl (B), 5% glycerol). Homogeneous monomeric polymerase was pooled and dialysed overnight with 6-8 kDa molecular weight cut-off membrane tubing (Spectra/Por, Spectrum Labs) into 50mM HEPES, 500 mM NaCl, 5% glycerol at pH 7. Finally, the protein was concentrated with Amicon® Ultra-15 (50 KDa cutoff), flash-frozen and stored at -80 °C.

### Negative stain electron microscopy

3 μl of protein solution were applied to glow-discharged carbon coated copper grid (300 mesh, Electron Microscopy Science) and let adsorb for 30 s. Grids were then washed twice in 25 μl drop of protein buffer and stained twice for 30 s with 6 μl of 2 % (w/v) uranyl acetate. Between each step, excess of protein/buffer solution/staining was blotted off using a filter paper. Grids were dried on adsorbing paper for at least 5 min. Negative-stain grids were imaged with a Tecnai 12 (FEI) TEM at 120 KV on a Ceta 16 M camera, at a nominal magnification of 16,000×.

### SEC-MALS analysis

Purified A/H7N9 polymerase was injected on Superdex200 10/300 SEC column (GE Healthcare) equilibrated in 50 mM HEPES (pH 7), 500 mM NaCl and 5% glycerol at 0.5 ml/min coupled to Wyatt Heleos II 18-angle light scattering instrument and to Wyatt Optilab rEX online refractive index detector (Wyatt Technology Corporation). Protein concentration was determined from the excess differential refractive index based on a 0.186 refractive index increment for a protein solution (1 g/ml). The concentration and the observed scattered intensity at each point in the chromatogram were used to calculate the absolute molecular mass from the intercept of the Debye plot using the Zimm model as implemented in Wyatt’s ASTRA software.

### Cap-dependent transcription assays

Separated 3′ and 5′ vRNA ends or loop mini-vRNA were used for transcription assays. 21-mer-sd or 21-mer-d 3′ and 14-mer 5′ ends vRNAs (IBA) were used in combination with synthetic 12-or 13-mer capped RNA (TriLink Biotechnologies) as primer (Table 2, Figure 1). A 56-mer loop vRNA was used in combination with a 6-mer primer. For the cap-dependent transcription assay, 0.2 μM A/H7N9 or FluB polymerase, 0.22 μM vRNAs, 0.4 μM capped RNA primer, 100 μM mix of NTPs, w/o 600 μM T1106-TP, and 2.5 pM α-^32^P-ATP were mixed and incubated in reaction buffer (150 mM NaCl, 50 mM HEPES, pH 7.4, 5 mM MgCl_2_ and 2 mM TCEP) at 28 °C for 4 hours.

**Table 2.**
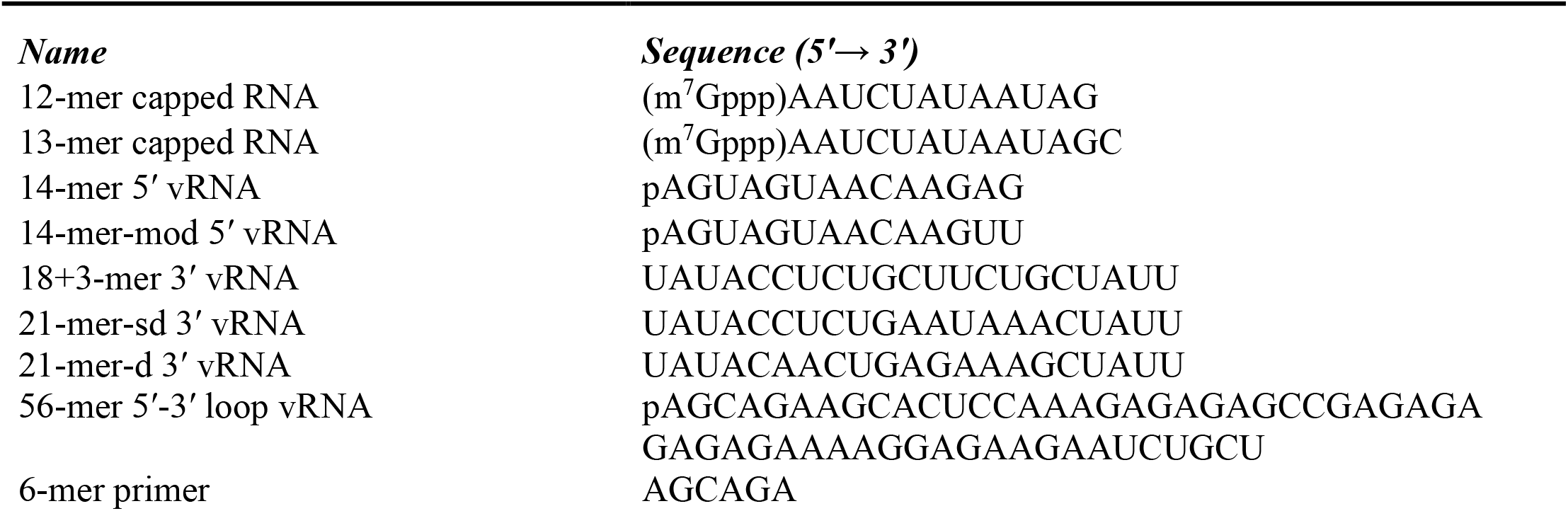
RNA sequences used for biochemical assays and cryoEM structures.

For the 6-mer dependent transcription assay, 1 μM FluB polymerase, 1.5 μM 56-mer vRNA, 20 μM 6-mer primer (Table 2), 100 μM NTPs, w/wo 1 mM 2′FdGTP or 600 μM T1106-TP and 0.04 μCi/μl α-^32^P-UTP or α-^32^P-ATP were mixed and incubated in reaction buffer (130 mM NaCl, 50 mM HEPES, pH 7.4, 5 mM MgCl_2_ and 0.4 mM TCEP) at 28 °C for 6 hours.

Samples were separated on a 7 M urea, 20% acrylamide gel in TBE buffer, exposed on a storage phosphor screen and read with a Typhoon scanner. The capped markers were synthetized as products of previously characterized transcription reactions (Kouba *et al*., 2019).

T1106-TP was synthesised as described (Huchting *et al*., 2018).

### CryoEM sample preparation and data collections

To capture distinct states of transcription cycle w/o T1106, specific sample preparation protocols (see below) were applied. All complexes were assembled in cryoEM buffer (50 mM HEPES pH 7.5, 150 mM NaCl, 5 mM MgCl_2_, 2 mM TCEP).

**Sample 1**: For the A/H7N9-Elongation complex, 0.8 μM FluA/H7N9 polymerase, 0.96 μM 3’ vRNA (18+3-mer) and 5’ vRNA (14-mer) and 1.6 μM 13-mer capped primer were mixed in cryoEM buffer and incubated for 4 hours at 28°C with 100 μM ATP and GTP, 300 μM CMPcPP, and then cooled to 4°C.

**Sample 2**: For the A/H7N9-Backtracked-T/U+1 complex, 0.9 μM A/H7N9 polymerase, 1.2 μM 3’ vRNA (21-mer-sd) and 5’ vRNA (14-mer) and 1.8 μM 12-mer capped primer were mixed in cryoEM buffer and incubated for 4 hours at 28°C with 100 μM UTP and CTP, and 600 μM T1106-TP and then cooled to 4°C.

**Sample 3**: For the B/Mem-Backtracked complex, 0.9 μM FluB polymerase, 1.2 μM 3’ vRNA (21-mer-d) and 5’ vRNA (14-mer-mod, Table 2) and 2 μM 13-mer capped primer were mixed in cryoEM buffer and incubated for 4 hours at 28°C with 100 μM UTP and CTP, and 600 μM T1106-TP and then cooled to 4°C.

**Sample 4:** For the A/H7N9-Apo dimer, 0.9 μM A/H7N9 polymerase was dialysed to 50 mM HEPES/NaOH, pH 7.5 and 570 mM NaCl.

**Sample 5:** For the A/H7N9-5’ hook-bound dimer and monomer, 0.9 μM A/H7N9 polymerase was first supplemented with 1.25 molar excess of 14-mer 5′ vRNA, 18+3-mer 3′ vRNA, and 15-mer capped primer and then dialysed for 12 hours to 50 mM HEPES/NaOH, pH 7.5 and 150 mM NaCl. Additional ~0.5 molar excess of 14-mer 5′ vRNA, 18+3-mer 3′ vRNA, and 15-mer capped primer was added to the sample mixture before plunge-freezing. Only the 14-mer 5′ vRNA was visible in the cryoEM structures.

For each sample, aliquots of 3 μl were applied to glow discharged grids (R2/1 or R1.2/1.3, Au 300, Quantifoil), blotted for 2 s and immediately plunge-frozen in liquid ethane using an FEI Vitrobot IV at 4°C and 100 % humidity. Grids were loaded onto 300 kV FEI Titan Krios and data were acquired in electron counting mode. Further details regarding each data collection are presented in Table 1 (elongation and backtracked structures) or Table S1 (A/H7N9 dimer structures).

### CryoEM image processing

All movie frames were aligned and dose-weighted using MotionCor2 program (Zheng *et al*, 2017). Thon rings, either from summed power spectra of every 4e^-^/Å^2^ or non-dose-weighted micrographs, were used for contrast transfer function parameter calculation with CTFFIND 4.1 (Zhang, 2016). Particles were selected with WARP (Tegunov & Cramer, 2019) or Gautomatch (http://www.mrc-lmb.cam.ac.uk/kzhang/). Further 2D and 3D cryo-EM image processing was performed in RELION 3.1 (Zivanov *et al*, 2020). First, particles were iteratively subjected to two rounds of 2D-classification and those in classes with poor structural features were removed.

#### 3D analysis of the of A/H7N9-Elongation complex

1.8 times binned particles (754.2 k) were subjected to global 3D auto-refinement with 60 Å low-pass filtered PDB ID 6T0V structure as initial model. The resulting global refinement was then subjected to global 3D classification (Figure EM-2) with fine angular searches using regularization parameter T = 4. The most defined class (116 k particles) was re-extracted to native pixel size and globally 3D auto-refined using aberration correction schemes in RELION.

#### 3D analysis of the A/H7N9-Backtracked-T/U+1 complex

1.8 times binned particles (1,067 k) were subjected to 3D-classifications with image alignment (Figure EM-4). The 3D-classification was restricted to eight classes and performed using 60 Å low-pass filtered PDB ID 6T0V structure as initial model. Particles in classes with poor structural features were removed. The remaining particles (776 k) were subjected to global 3D auto-refinement and then subjected to another round of global 3D classification with fine angular searches using regularization parameter T = 4. The most defined class (172 k particles) was re-extracted to native pixel size and globally 3D auto-refined. The resulting global refinement was subjected to final round of 3D-classification focused on the core region of the complex. The most defined class (98.9 k particles) was globally 3D auto-refined using aberration correction schemes in RELION.

#### 3D analysis of the of B/Mem-Backtracked complex

3.52 times binned particles (5,479 k) were globally 3D auto-refined and then subjected to 3D-classifications with image alignment restricted to eight classes (Figure EM-6). Particles in classes with poor structural features were removed and remaining particles (3,619 k) were re-extracted and subjected to another round of global 3D auto-refinement followed by 3D-classifications to ten classes. Particles in classes with poor structural features were removed and remaining particles (1,540 k) were re-extracted to native pixel size, globally 3D auto-refined and finally focus-refined on the core region of the complex. The resulting global refinement was subjected to final round of 3D-classification with fine angular searches using regularization parameter T = 12, restricted to 8 classed and focused on the core region of the complex. The most defined classes were pooled (328,2 k particles) and globally 3D auto-refined using aberration correction schemas in RELION.

#### 3D analysis of the of A/H7N9 apo-dimer

2 times binned particles (314.8 k) were subjected to global 3D classification with 60 Å low-pass filtered PDB ID 3J9B structure as initial model. The most defined classes (156.7) were pooled, re-extracted to native pixel size, and globally 3D auto-refined. The resulting global refinement was then subjected to global 3D classification (Figure EM-8) with fine angular searches using regularization parameter T = 8. The most defined class (101.9 k particles) was globally 3D auto-refined using C2 symmetry and aberration correction schemes in RELION.

#### 3D analysis of the of A/H7N9 5’ hook bound dimer

Un-binned particles (84.2 k) were subjected to global 3D classification (Figure EM-10) with 60 Å low-pass filtered PDB ID 3J9B structure as initial model using regularization parameter T = 4. The most defined class (58.8 k particles) was globally 3D auto-refined using C2 symmetry and aberration correction schemes in RELION.

#### 3D analysis of the of A/H7N9 5’ hook bound monomer

2 times binned particles (500.9 k) were subjected to global 3D classification with 60 Å low-pass filtered one protomer PDB ID 3J9B structure as initial model. The most defined classes (124 k) were pooled, re-extracted to native pixel size, and globally 3D auto-refined. The resulting global refinement was then subjected to global 3D classification (Figure EM-10) with fine angular searches using regularization parameter T = 4. The most defined class (37.7 k particles) was globally 3D auto-refined using aberration correction schemes in RELION.

All final cryo-EM density maps were generated by the post-processing feature in RELION and sharpened or blurred into MTZ format using CCP-EM (Burnley *et al*, 2017). The resolutions of the cryo-EM density maps were estimated at the 0.143 gold standard Fourier Shell Correlation (FSC) cut off (Figure EM 1d, 3d, 5d, 7d, 9d). A local resolution (Figure EM 1e, 3e, 5e, 7e, 9e) was calculated using RELION and reference-based local amplitude scaling was performed by LocScale (Jakobi *et al*, 2017).

### CryoEM model building and refinement

The A/H7N9 elongation and backtracked and FluB backtracked structures were constructed by first rigid-body fitting into the cryoEM density using respectively the promoter bound A/H5N1 (PDB:6RR7) and FluB elongation (PDB:6QCT) polymerase structures as starting points. For the apo- and 5′ hook bound A/H7N9 dimer structures a previous crystal structure of a truncated A/H7N9 dimer (PDB:6TU5) was used as initial model. The models were iteratively improved by manual adjusting with COOT(Emsley & Cowtan, 2004) and refinement with PHENIX real-space-refinement (Afonine *et al*, 2018). Validation was performed using the PHENIX validation tool and model resolution was estimated at the 0.5 FSC cut off (Table 1, Table S1).

### Computational Methods

#### Models systems

The experimental A/H7N9 backtracked structure was used to set up a model system of the polymerase. Missing loops in the protein structure were added using SwissModel (Waterhouse *et al*, 2018). The protein/RNA complex was solvated in a rhombic dodecahedral box of water molecules, with a buffer distance of 16 Å between each wall and the closest atom in each direction. The system was then neutralized with K+, and additional K+ and Cl− ions were added to reach ∼100 mM ionic concentrations. The final model includes a total of ∼241,000 atoms. To investigate the conformational properties of the U:T1106 base pair incorporated in an RNA double helix, two RNA dodecamers with different sequences were considered (T = T1106):

CCCUCUGCUUGG

GGGAGTCGAACC

in which the U:T1106 pair is flanked by CG pairs, and

CGCGAAUUCGCG

GCGCUUTAGCGC

in which the U:T1106 pair is flanked by AU pairs. Duplexes structures were built in standard helical A-form. For each sequence, initial geometries corresponding to canonical and wobble U:T1106 base pairing were generated superimposing the Density Functional Theory (DFT)-optimized geometries to the corresponding base pair. Each duplex was solvated with ~9000 water molecules and 22 sodium ions were added randomly to neutralize the charge of the systems.

#### Quantum Mechanical calculations

Quantum chemical calculations were performed using the hybrid B3LYP (Becke, 1993; Lee *et al*, 1988) density functional method as implemented in Gaussian09 (https://gaussian.com/glossary/g09/). Geometry optimization was performed using the 6-31G* basis set, accounting for solvent effects (water) via the PCM continuum model (Tomasi *et al*, 2005).

#### MD simulations

Molecular dynamics (MD) simulations were performed with the pmemd module of Amber20 (https://ambermd.org/doc12/Amber20.pdf). The AMBER-ff14SB (Maier *et al*, 2015) and AMBER-ff12SB (ff99 + bsc0 + χOL3) (Perez *et al*, 2007) force fields were used for protein and RNA, respectively. Atom types and bonding parameters for the m^7^G cap and the T1106 drug were adapted from the same AMBER force field, and atomic point charges were derived using the RESP procedure (Bayly *et al*, 1993). Monovalent and divalent metal ions were described with Li and Merz 12-6 parameters (Li *et al*, 2015). The TIP3P model was adopted for water (Jorgensen *et al*, 1983). Simulations were performed with a distance cutoff of 10 Å. Long-range electrostatics were treated with the particle mesh Ewald method (Li *et al*, 2013). Bonds involving hydrogen atoms were constrained, allowing a time step of 2 fs. After solvent equilibration, the entire system was energy minimized and gently heated to 310 K during 0.5 ns while restraining protein and RNA backbone atoms. The Andersen-like temperature-coupling scheme (Åqvist *et al*, 2004) and a Monte Carlo barostat (Åqvist *et al*., 2004) were used to maintain temperature and pressure close to room temperature conditions. About 1 μs MD simulation in the NPT ensemble was accumulated for each model system.

#### Structure data availability

Influenza A/H7N9 polymerase elongation complex PDB:7QTL and EMDB:EMD-14144 Early transcription elongation state of influenza A/H7N9 polymerase backtracked due to double incorporation of nucleotide analogue T1106 and with singly incorporated T1106 at the +1 position PDB:7R0E and EMDB:EMD-14222

Early transcription elongation state of influenza B polymerase backtracked due to double incorporation of nucleotide analogue T1106 PDB:7R1F and EMDB:EMD-14240 Influenza A/H7N9 apo-dimer complex PDB: 7ZPM and EMDB:EMD-14858

Influenza A/H7N9 5’ hook bound dimer complex PDB: 7ZPL and EMDB:EMD-14857

## Notes

### Competing Interest Statement

The authors have declared no competing interest.

