## Supplemental Table S1, Supplemental Figures and Legends, Supplemental CryoEM Methods for "Backtracking of influenza polymerase upon consecutive incorporation of nucleoside analogue T1106 directly observed by high-resolution cryo-electron microscopy"

#### Supplementary Material

**Contents:**

**Supplementary Table S1**

**Supplementary Figure Legends**

**Supplementary Figures**

**CryoEM structure determination details.**

**Table S1. CryoEM structure determination and validation statistics of dimeric A/H7N9 structures**

| Name of structure | A/H7N9 apo-dimer | A/H7N9 5' hook dimer |
|---|---|---|
| PDB ID | 7ZPM | 7ZPL |
| EMDB ID | EMD-14858 | EMD-14858 |
| **Data collection and processing** | | |
| Microscope | FEI Titan Krios | FEI Titan Krios |
| Voltage (kV) | 300 | 300 |
| Camera | Gatan K2 Quantum | Gatan K2 Quantum |
| Magnification | 165,000 | 165,000 |
| Nominal defocus range (μm) | 0.5 – 3.0 | 0.4 – 3.0 |
| Exposure time (s) | 6 | 4 |
| Electron exposure (e–/Å$^2$) | 41.4 | 31.2 |
| Number of frames collected (no.) | 40 | 32 |
| Number of frames processed (no.) | 40 | 32 |
| Pixel size (Å) | 0.831 | 0.84 |
| Micrographs (no.) | 4048 | 5,381 |
| Total particle images (no.) | 821738 | 1,096,109 |
| **Refinement** | | |
| Particles per class (no.) | 101,870 | 58,775 |
| Map resolution (Å), 0.143 FSC | 2.81 | 3.12 |
| Model resolution (Å), 0.5 FSC | | |
| Map sharpening *B* factor (Å$^2$) | -73.78 | -21.9 |
| Map versus model cross-correlation (CCmask) | 0.8830 | 0.8824 |
| **Model composition** | | |
| Non-hydrogen atoms | 19330 | 19919 |
| Protein residues | 2412 | 2422 |
| Nucleotide residues | - | 24 |
| Water | - | - |
| Ligands | - | 1 (Mg) |
| ***B* factors (Å$^2$)** | | |
| Protein | 71.75 | 92.96 |
| Nucleotide | - | 69.26 |
| Ligand | - | 80.78 |
| Water | - | - |
| **R.M.S. deviations** | | |
| Bond lengths (Å) | 0.002 | 0.002 |
| Bond angles (°) | 0.496 | 0.389 |
| **Validation** | | |
| MolProbity score | 1.62 | 1.35 |
| All-atom clashscore | 6.91 | 5.65 |
| Poor rotamers (%) | 0.14 | 0.0 |
| **Ramachandran plot** | | |
| Favoured (%) | 96.40 | 97.84 |
| Allowed (%) | 3.52 | 2.16 |
| Outliers (%) | 0.08 | 0 |

#### Supplementary Figure Legends

##### Figure S1. Purification of A/H7N9

**A**) Analysis of the molecular mass of the purified A/H7N9 apo-polymerase by size-exclusion chromatography (SEC) coupled to MALS (multi-angle light scattering). The analysis (black line) indicates an average molecular mass of the complex between 400 and 500 kDa across the main peak of the chromatogram (black line). The expected molecular mass of the A/H7N9 polymerase dimer is ~510 kDa.

**B**) SEC analysis of A/H7N9 apo-polymerase (orange line) and A/H7N9 polymerase bound to vRNA promoter (black line). When the vRNA promoter is bound, the A/H7N9 polymerase migrates in SEC analysis primarily as a monomer.

**C**) SDS-PAGE analysis of the purified A/H7N9 apo-polymerase. 40 μg protein sample fractions after SEC-MALS analysis are indicated by the elution volume. The first lane contains the molecular weight marker.

D) Negative-stain electron microscopy images of the A/H7N9 apo-polymerase (**left**) and A/H7N9 polymerase bound to vRNA promoter (**right**). Dimer and monomer particles are marked in orange and white circles, respectively.

##### Figure S2. CryoEM density for the RNA in the three determined structures.

**A**) A/H7N9 transcription structure stalled by CMPcPP at overall 2.48 Å resolution.

**B**) T1106-stalled and backtracked A/H7N9 structure at overall 2.51 Å resolution.

**C**) T1106-stalled and backtracked FluB structure at overall 2.58 Å resolution.

In each case, a stick model of the entire RNA (template yellow, capped product slate blue and 5' end violet) is shown superposed with the cryoEM density (top) together with a detail of (**A**) the G:CMPcPP base-pair, (**B**) the U:T1106 wobble base-pair and (**C**) a G:C base-pair, each in the +1 position (bottom).

##### Figure S3: CryoEM structure of A/H7N9 apo- and 5' hook-bound dimers.

**A**) Ribbon diagrams of the overall A/H7N9 apo- (top) and 5' hook-bound (bottom) symmetrical dimer structures looking down the 2-fold axis. One polymerase trimer is coloured in each case with PA (green), PB1 (cyan), PB2 (red) or the corresponding light colours for the 5' hook or apo-dimers, respectively. The other polymerase trimer is grey. The 5' hook is violet.

**B**) Comparison of the A/H7N9 apo- (left) and 5' hook-bound (right) polymerase trimers looking into the active site cavity, coloured as in **A**. The change in conformation of the PB1/23-35 peptide and structuring of the fingertips loop (violet-blue) upon 5' hook binding are indicated.

**C**) CryoEM density for the PB1/23-35 and 503-513 peptides in the A/H7N9 apo-dimer (left) and 5' hook-bound (right) structures.

**D**) Superposition of the A/H7N9 apo- and 5' hook-bound (bottom) symmetrical dimer structures showing concerted conformational changes that occur upon 5' hook-binding. Colours as in **A**.

**Figure S4. Comparison of the conformation and interactions of capped primer moiety in various structures.**

**A**) Conformation of the capped product (slate blue) in the A/H7N9 elongation structure. Interacting residues from PB2 are shown and coloured red, magenta and orange if they are from the PB2-N, midlink and cap-binding domains respectively.

**B**) Comparison of the conformation of the cap-proximal part of the capped primer/product (slate blue) in (from left to right) the A/H7N9 elongation structure (PDB:7QTL), bat A/H17N10 elongation structure, in which only three bases are clearly visible in the cryoEM structure (PDB:6T0V), FluB/Memphis elongation structure (PDB:6QCT) and FluB/Memphis backtracked structure (PDB:7R1F). Key residues (see text) from PB2 are shown and coloured magenta and orange for the midlink and cap-binding domains respectively.

**C**) Comparison of the RNA from the A/H7N9 elongation (capped product: slate blue, template: orange) and backtracked (capped product: light blue, template: light orange) structures. Upon backtracking, there is shorter connection from the cap to the product-template duplex (7 nucleotides compared to 9 in the elongation structure), eliminating the looped out region in the elongation structure.

**Figure S5. Molecular dynamics on the T1106:U base-pair in different environments.**

**A**) Relative energies of the DFT optimized structures of the canonical (left) and wobble (right) base pairing between U and T1106 (R=H) or T705 (R=F). Energies in kcal/mol.

**B**) Time evolution of the root mean square deviation (RMSD) for the heavy atoms of nucleotide U(6) and T1106, using as reference the backtracked A/H7N9 crystal structure. (Right) Schematic representation of nucleotide U(6) and T1106.

**C**) Time evolution of the distance between the nitrogen atom of the K229 side chain (NK229) and the oxygen of the T1106 amide (OT1106). (Right) Schematic representation of RNA double helix portion in correspondence of T1106:U(6) base pair, and K229 residue. The dashed line corresponds to the distance between K229 side chain and T1106 amide.

**D**) (Left) Time evolution during the MD simulation of the root mean square deviation (RMSD) for the heavy atoms of the influenza A/H7N9 polymerase backtracked structure, using as reference the cryoEM structure. (Right) Schematic representation of RNA-RdRp A/H7N9 backtracked structure.

**Figure S6. Stalling during processive elongation.**

**A**) 56-mer-d template (yellow) with T1106 single double-incorporation site (CC) at position (31-32). Expected products from transcription reactions using 6-mer primer (blue) and combinations of nucleotides and 2FdGTP or T1106-TP.

**B**) Gel showing products of the reactions schematised in (**A**).

### Figure S1. Characterisation of A/H7N9 polymerase

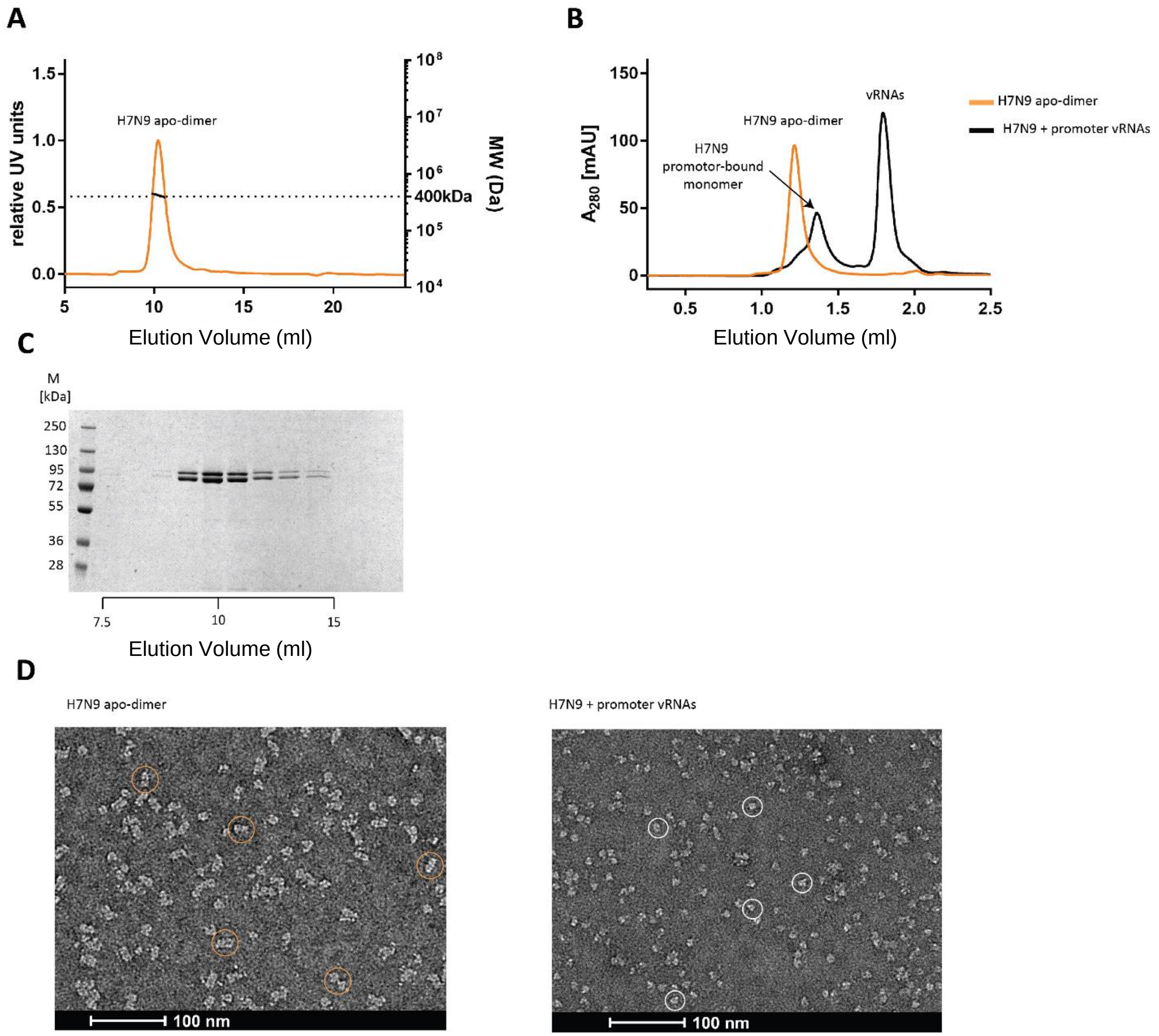

**Figure S2.** **Cryo EM density for elongation and backtracked structures**

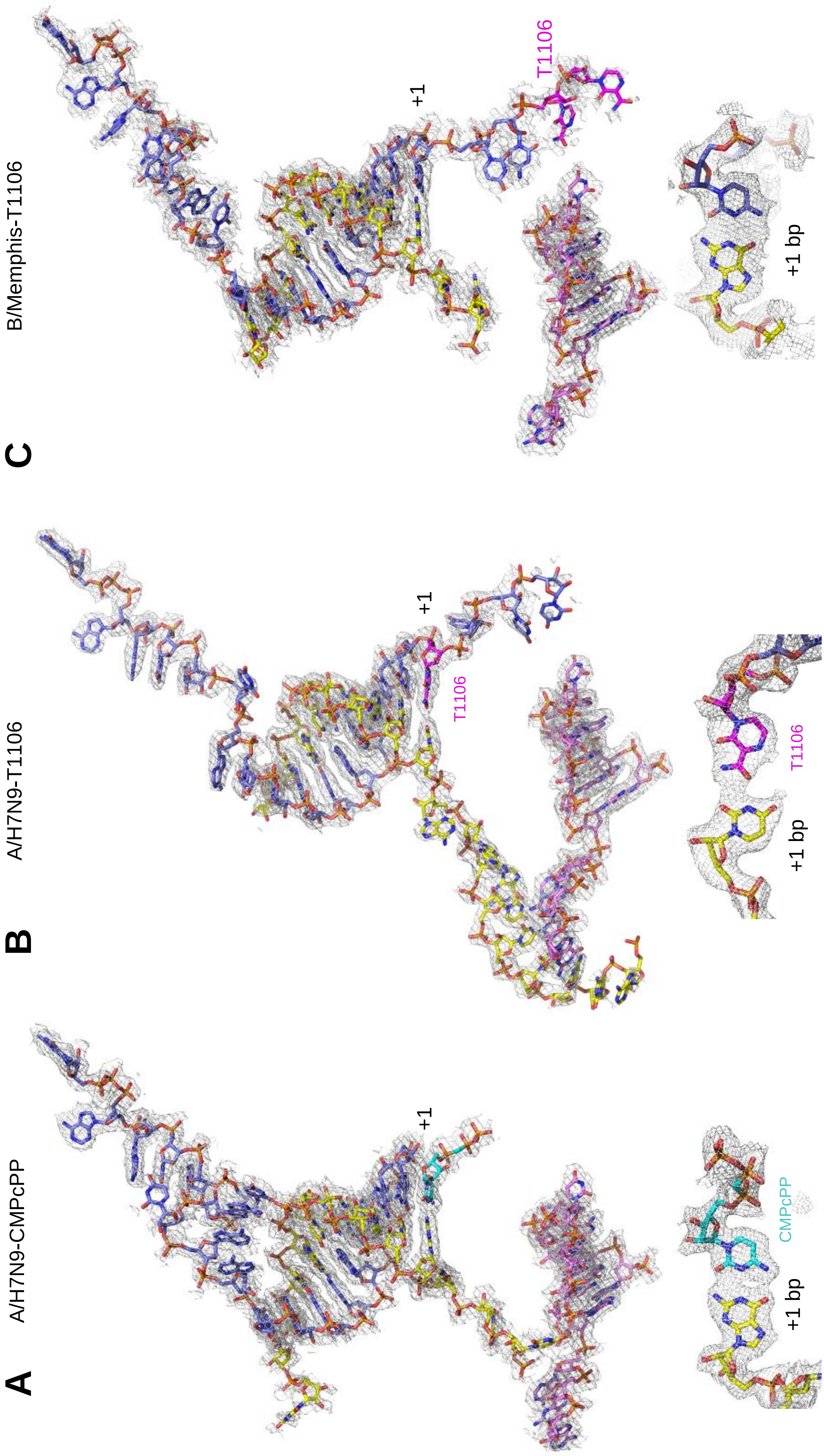

### Figure S3. A/H7N9 apo- and 5' hook-bound dimers

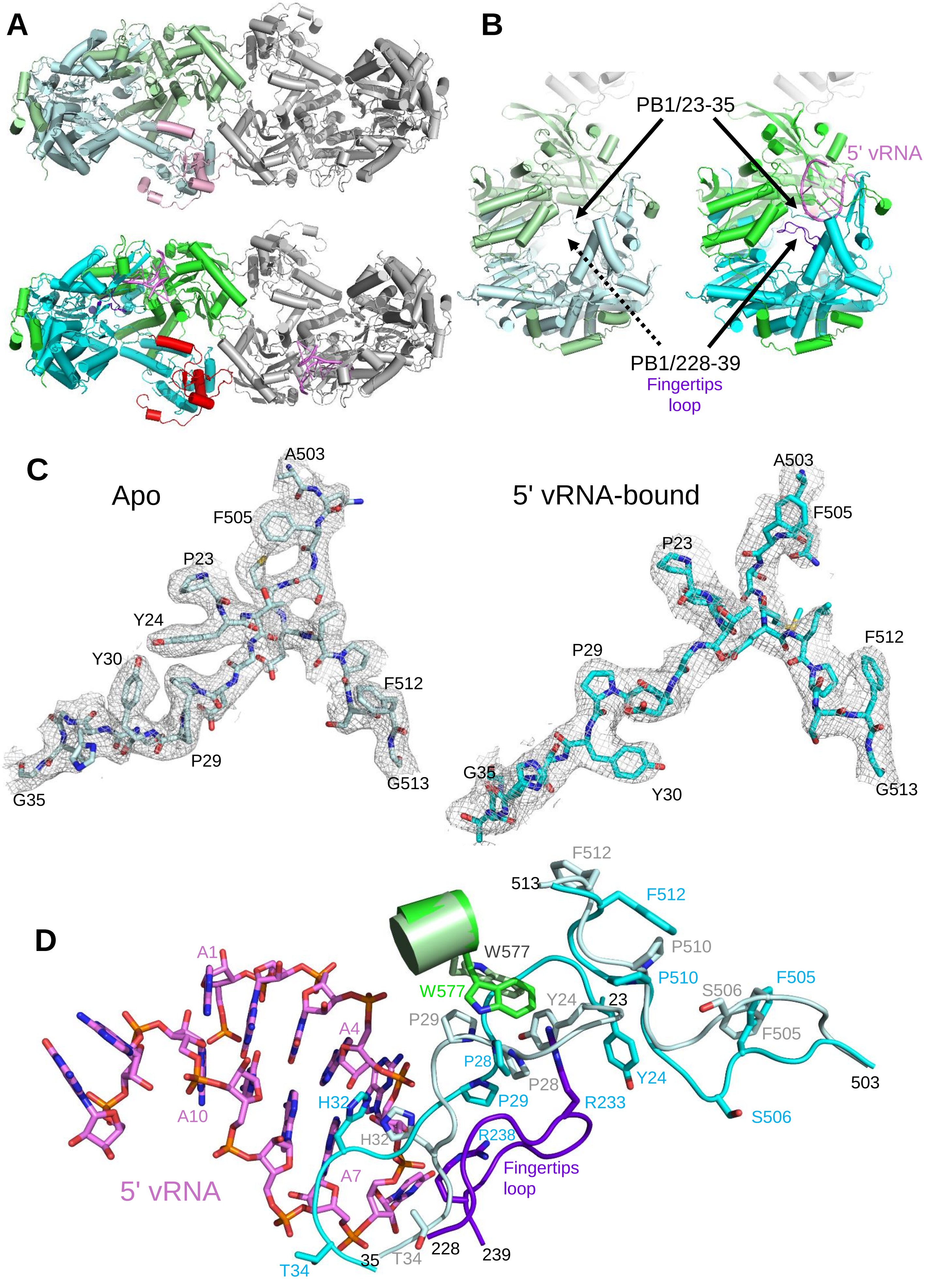

### Figure S4. Conformation and interactions of capped primer moiety

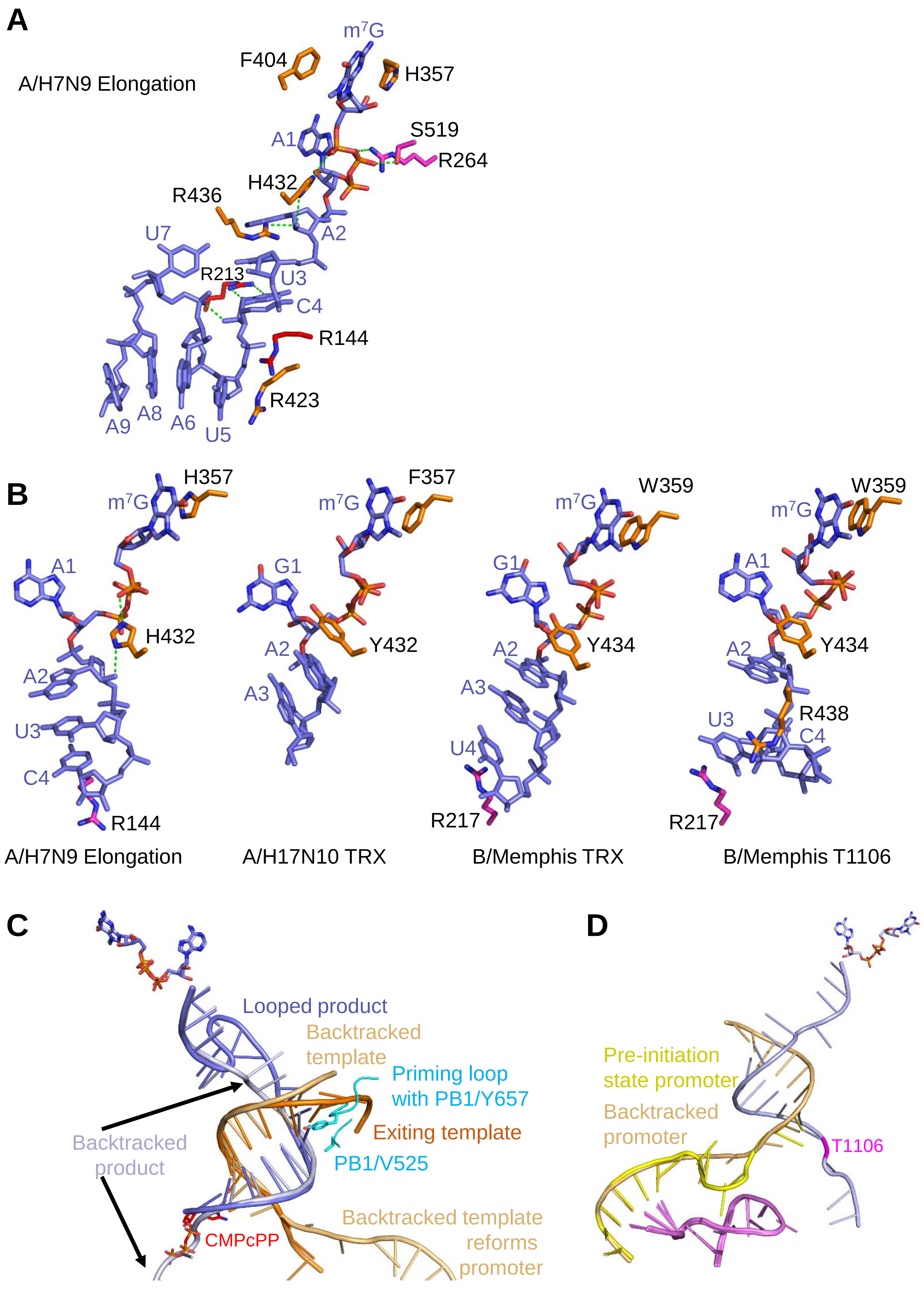

**Figure S5. Molecular dynamics of the T1106:U base-pair**

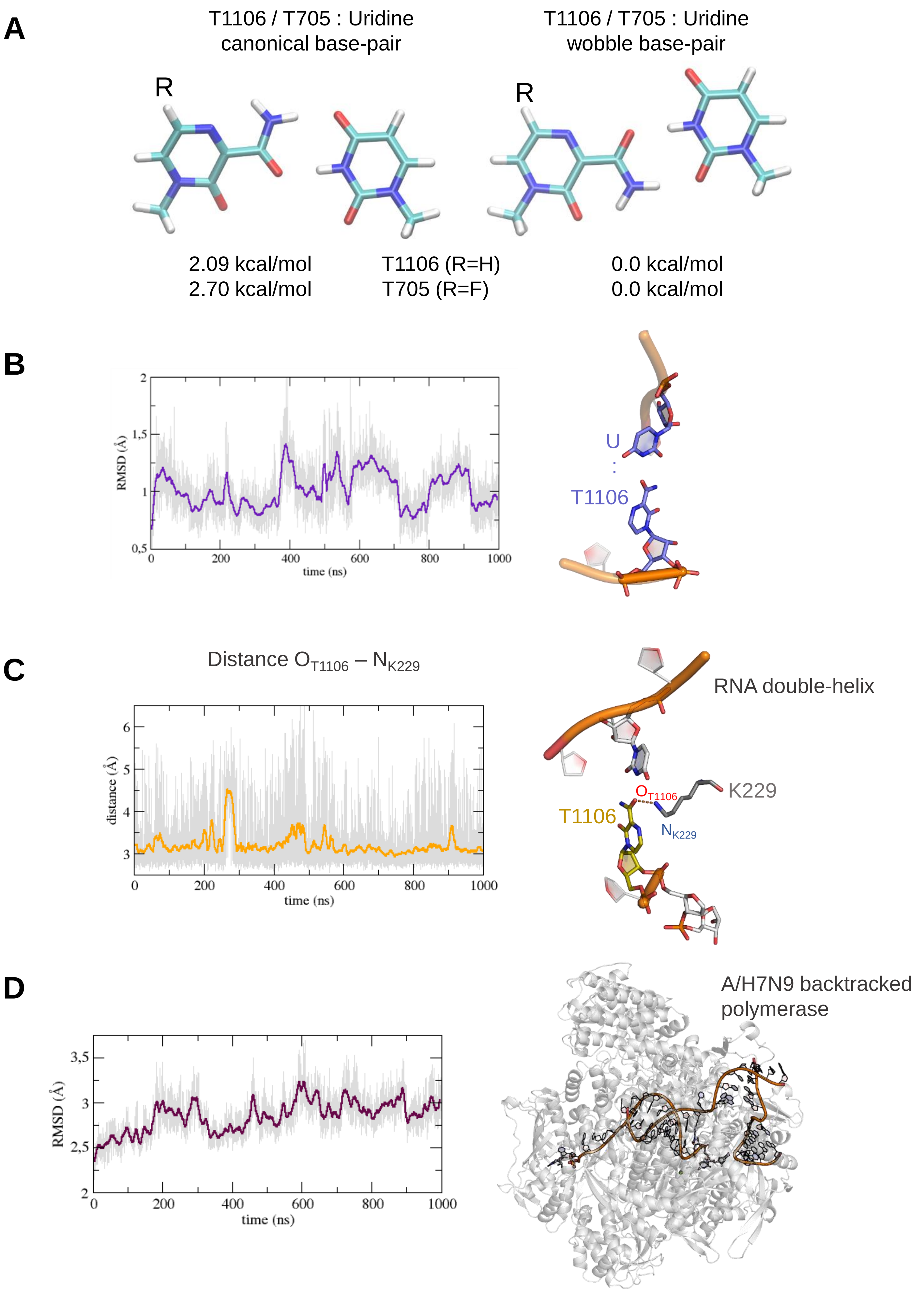

### Figure S6. Stalling during processive elongation

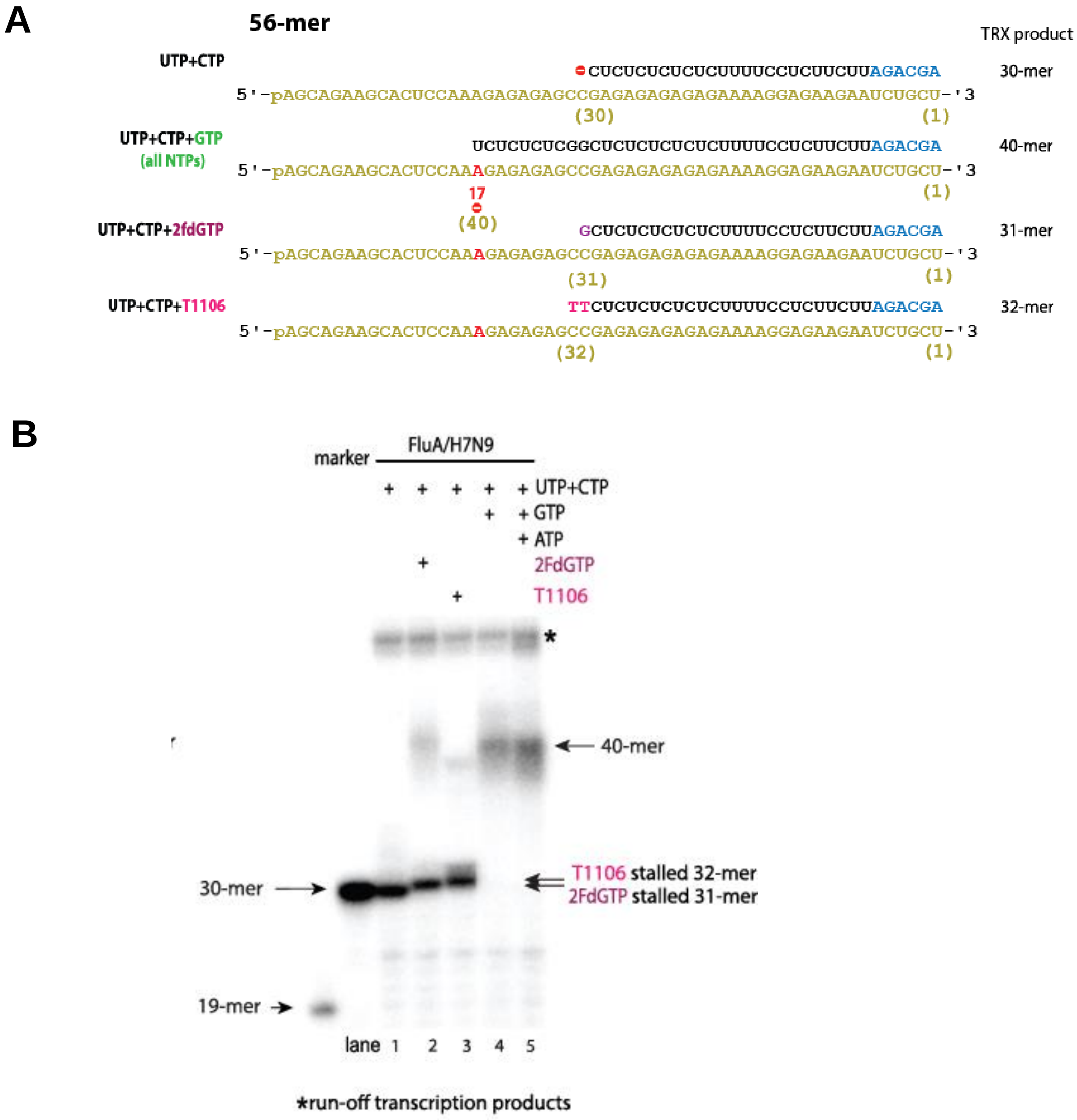

#### CryoEM structure determination details.

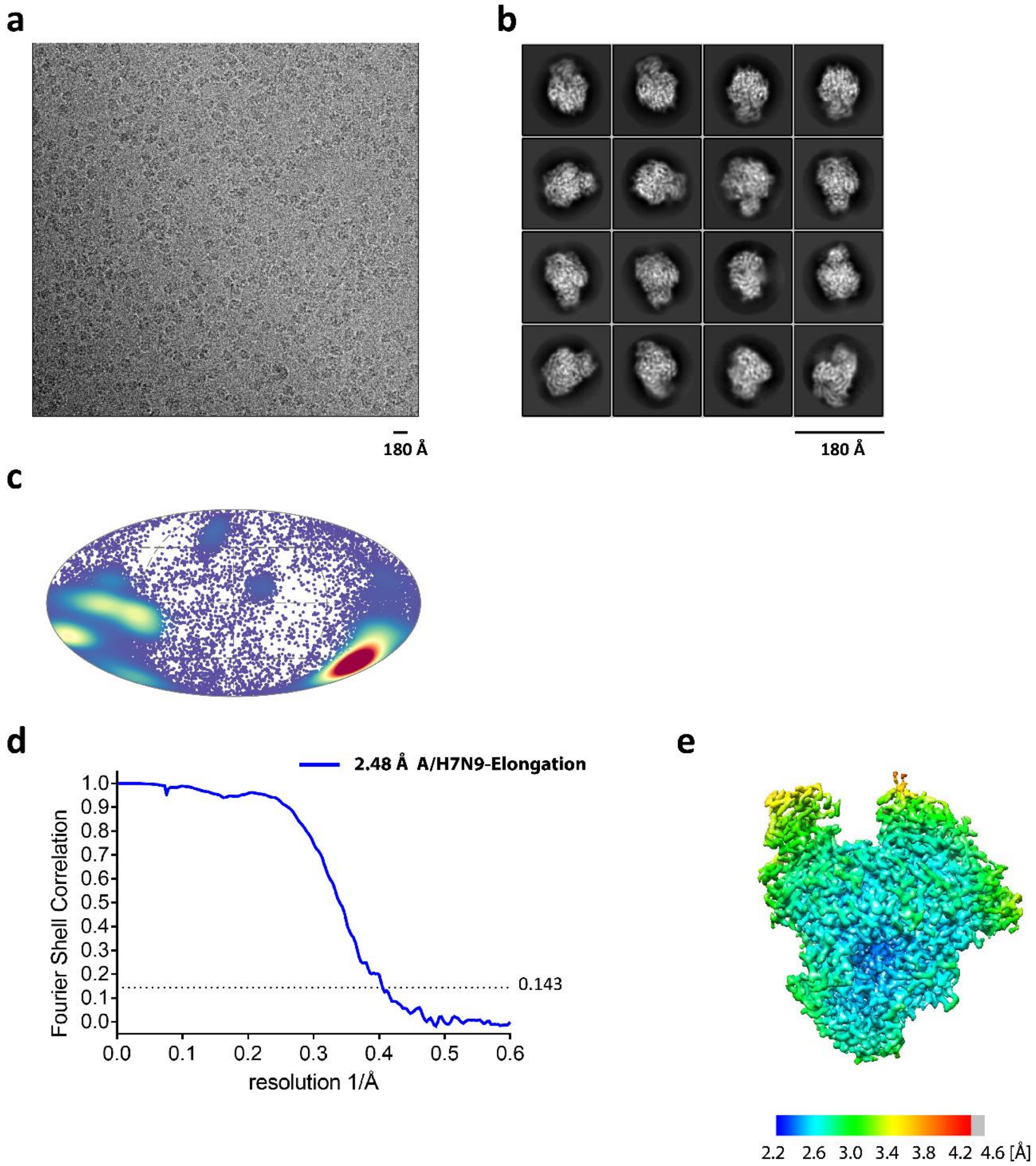


**Figure EM-1: Cryo-EM of A/H7N9-Elongation complex.**

**a**, Micrograph of the A/H7N9-Elongation complex in free standing ice after MotionCor2 (Zheng et al., 2017) correction at defocus of ~2.5 µm. **b**, 2D-class averages of the A/H7N9-Elongation complex **c**, Angular distribution for particles of the A/H7N9-Elongation complex visualized on a globe-like plane. **d**, Fourier shell correlation (FSC) curves for the A/H7N9-Elongation complex (blue). The plot of the FSC between two independently refined half-maps shows the overall resolution of the two maps as indicated by the gold standard FSC 0.143 cut-off criteria (Rosenthal and Henderson, 2003). **e**, Surface representation of local resolution distribution of the A/H7N9-Elongation complex. The map is colored according to the local resolution (as indicated in the color bar) calculated within the RELION software package.

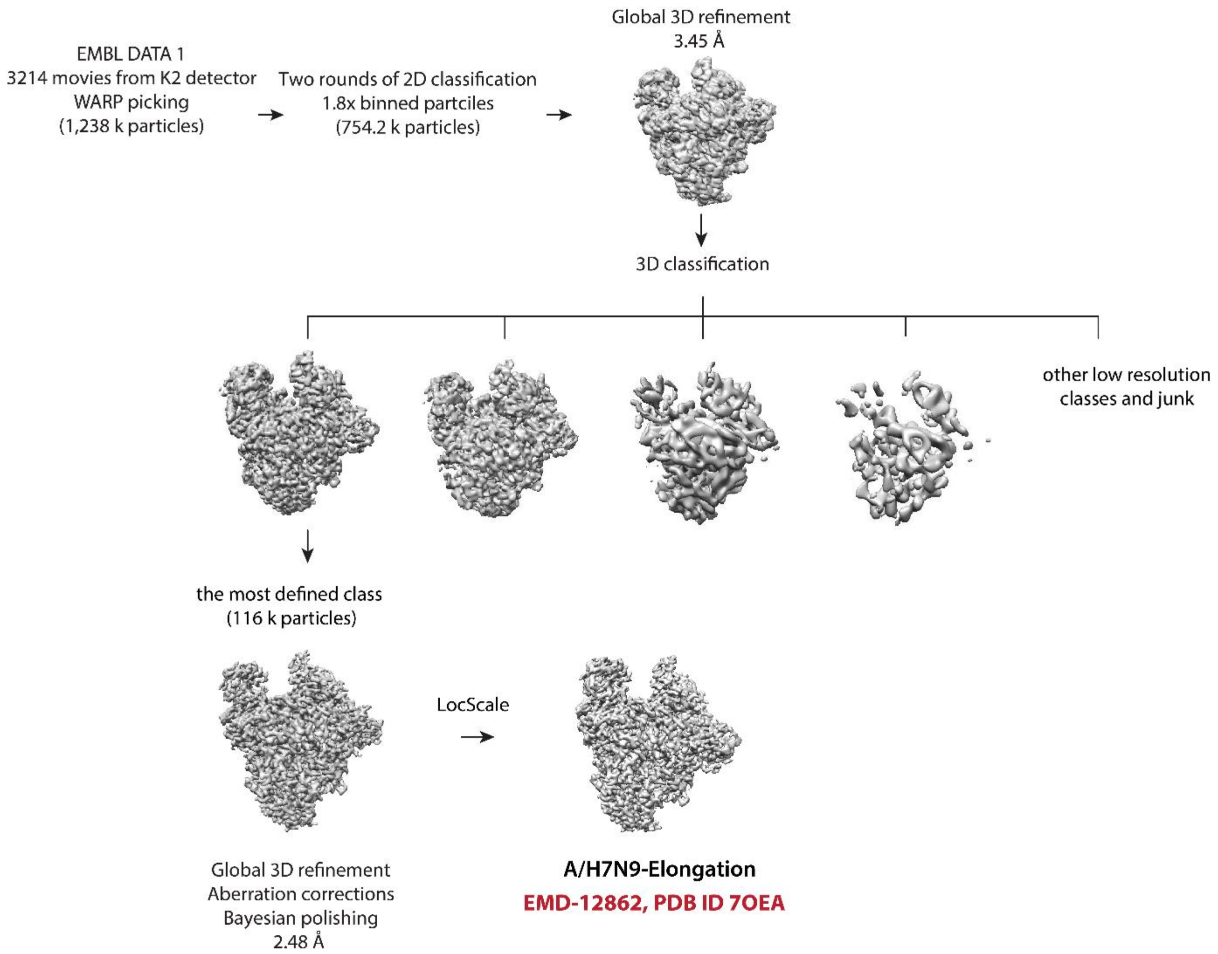


**Figure EM-2: Cryo-EM data 3D classification and refinement scheme of A/H7N9-Elongation complex.**

Summary of the cryo-EM 3D classification and refinement scheme of the A/H7N9-Elongation complex. EMBL DATA 1 2D classes with well-defined secondary structure features were merged (754.2 k particles). The merged particles were globally 3D-refined and then classified into six 3D classes with angular assignment. Incomplete, low resolution, and damaged particle classes were excluded from further data analyses. The best-defined A/H7N9-Elongation complex class was globally 3D-refined using aberration correction schemes. The final cryo-EM map was refined and post-processed with a respective mask in RELION 3.1 (Zivanov et al., 2018; Zivanov et al., 2020) and filtered by LocScale (Jakobi et al., 2017).

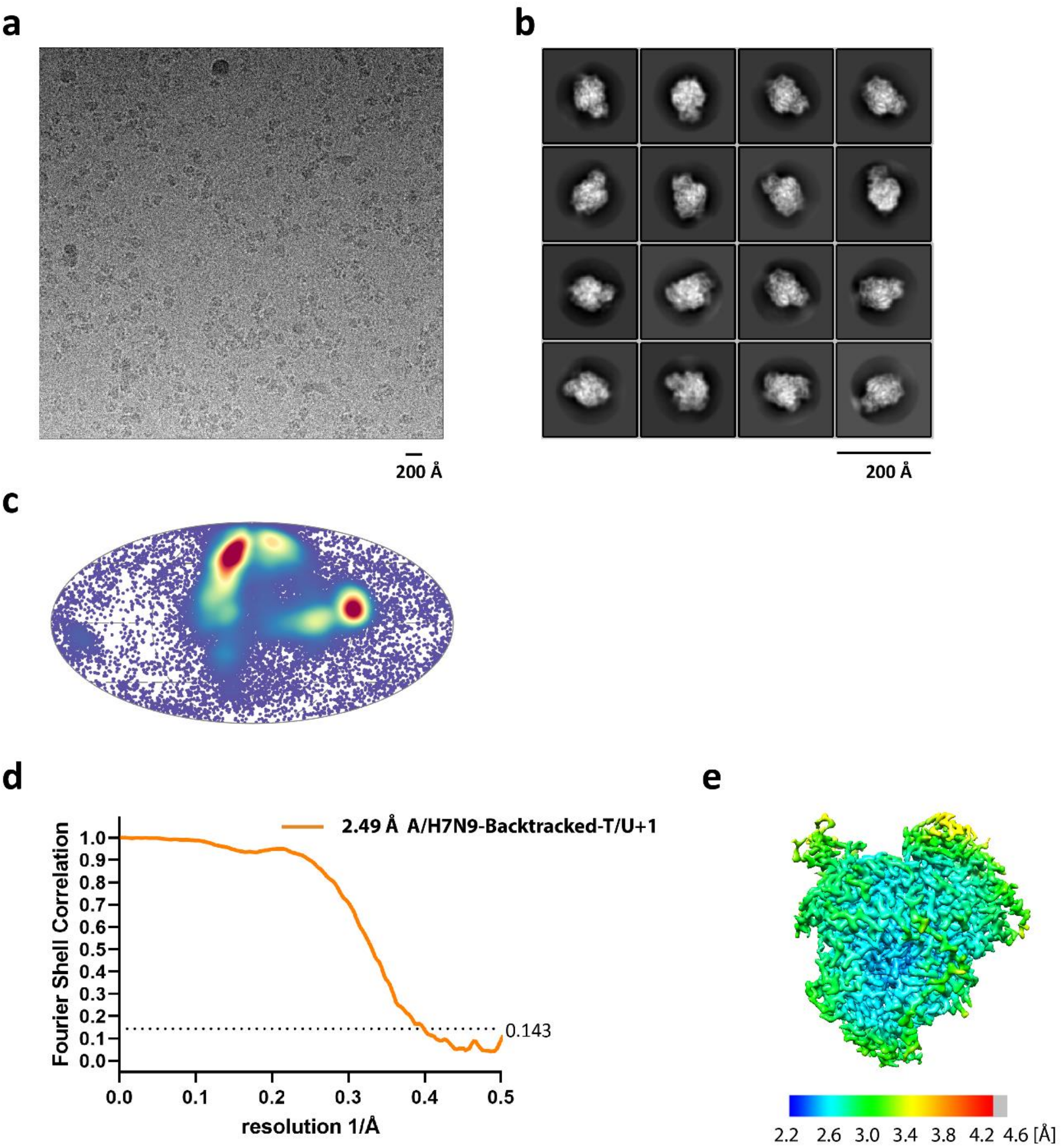


**Figure EM-3: Cryo-EM of A/H7N9-Backtracked-T/U+1 complex.**

**a**, Micrograph of the A/H7N9-Backtracked-T/U+1 complex in free standing ice after MotionCor2(Zheng et al., 2017) correction at defocus of ~2.5 µm. **b**, 2D-class averages of the A/H7N9-Backtracked-T/U+1 complex. **c**, Angular distribution of particle projections of the A/H7N9-Backtracked-T/U+1 complex visualized on a globe-like plane. **d**, Fourier shell correlation (FSC) curves for the A/H7N9-Backtracked-T/U+1 complex (orange). The plot of the FSC between two independently refined half-maps shows the overall resolution of the two maps as indicated by the gold standard FSC 0.143 cut-off criteria (Rosenthal and Henderson, 2003). **e**, Surface representation of local resolution distribution of the A/H7N9-Backtracked-T/U+1 complex. The map is colored according to the local resolution (as indicated in the color bar) calculated within the RELION software package.

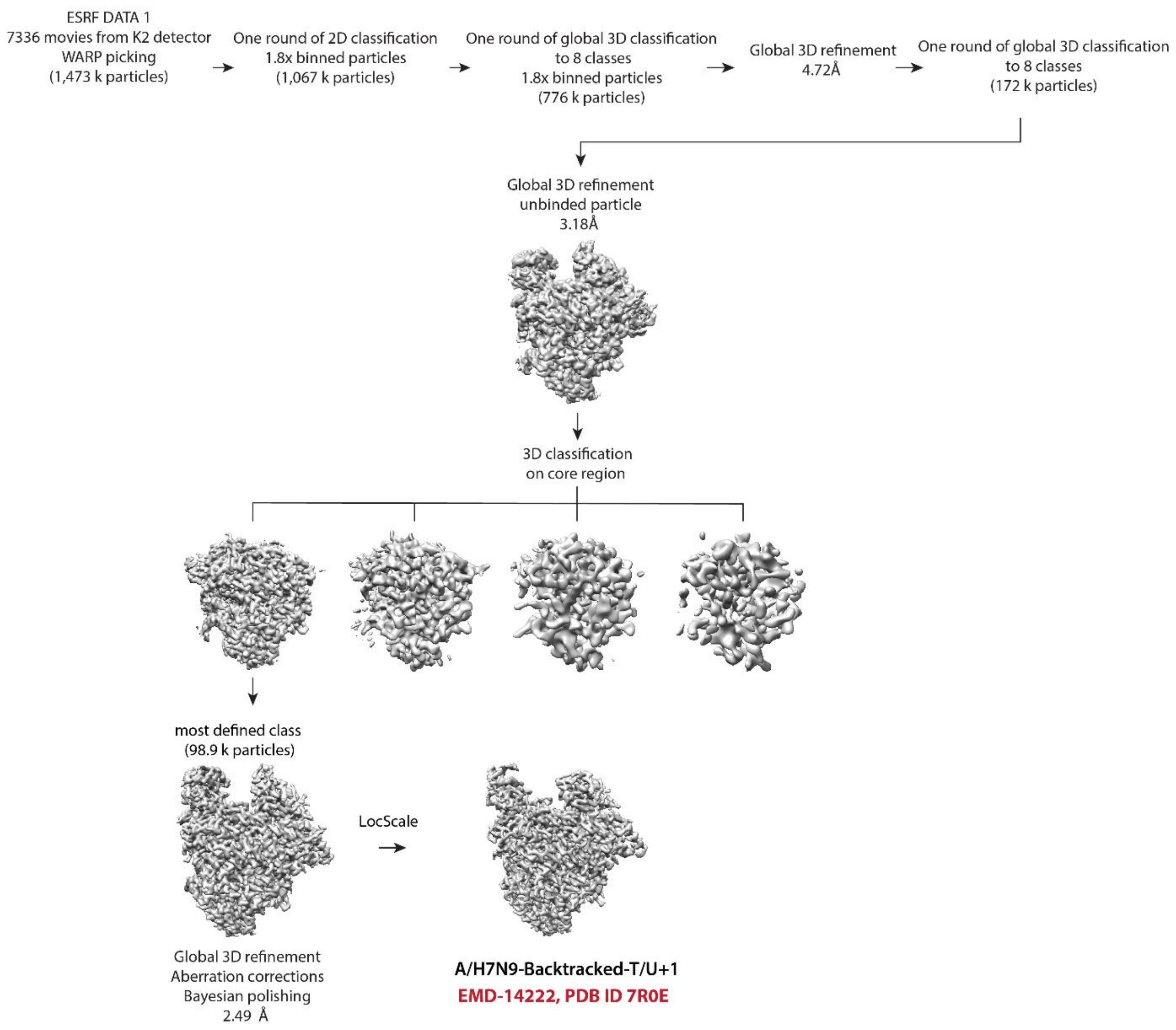


**Figure EM-4: Cryo-EM data 3D classification and refinement scheme of A/H7N9-Backtracked-T/U+1 complex.**

Summary of the cryo-EM 3D classification and refinement scheme of the A/H7N9-Backtracked-T/U+1 complex. ESRF DATA 1 2D classes with well-defined secondary structure features were merged (1,067 k particles). The merged particles were globally 3D-classified and incomplete, low resolution, and damaged particle classes were excluded from further data analyses. The remaining particles (776 k) were globally 3D-refined and subsequently 3D-classified. The most defined classes was merged (172 k), re-extracted to native pixel, globally 3D-refined and finally focus 3D-classified around the core region of the complex. The most defined class was globally 3D-refined using aberration correction schemes. The final cryo-EM map was refined and post-processed with a respective mask in RELION 3.1 (Zivanov et al., 2018; Zivanov et al., 2020) and filtered by LocScale (Jakobi et al., 2017).

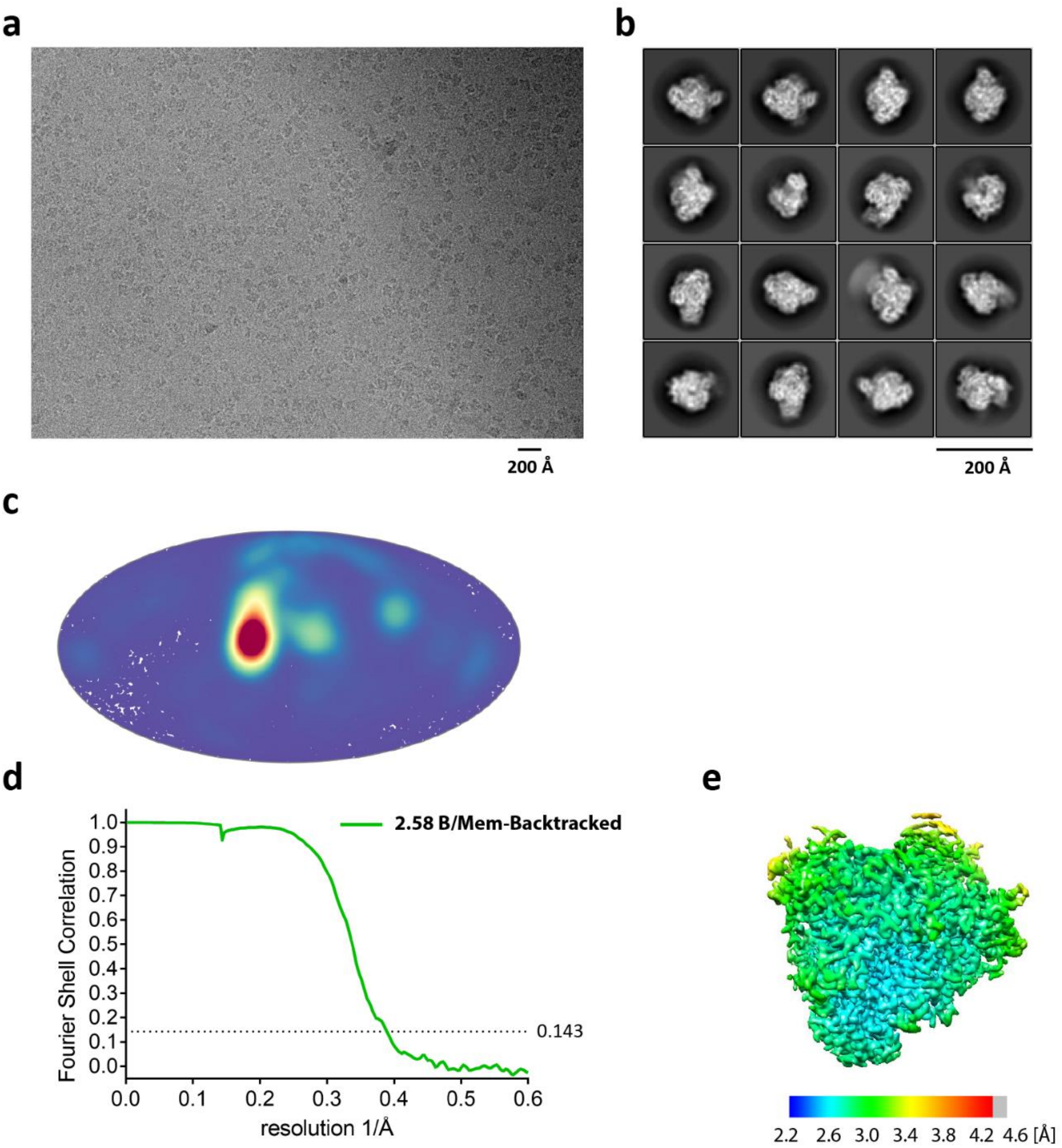


**Figure EM-5: Cryo-EM of B/Mem-Backtracked complex**

**a**, Micrograph of the B/Mem-Backtracked complex in free standing ice after MotionCor2(Zheng et al., 2017) correction at defocus of ~2.5 µm. **b**, 2D-class averages of the of the B/Mem-Backtracked complex **c**, Angular distribution for particle projections of the B/Mem-Backtracked complex visualized on a globe-like plane. **d**, Fourier shell correlation (FSC) curves for the B/Mem-Backtracked complex. The plot of the FSC between two independently refined half-maps shows the overall resolution of the two maps as indicated by the gold standard FSC 0.143 cut-off criteria (Rosenthal and Henderson, 2003). **e**, Surface representation of local resolution distribution of the B/Mem-Backtracked complex. The map is colored according to the local resolution (as indicated in the color bar) calculated within the RELION software package.

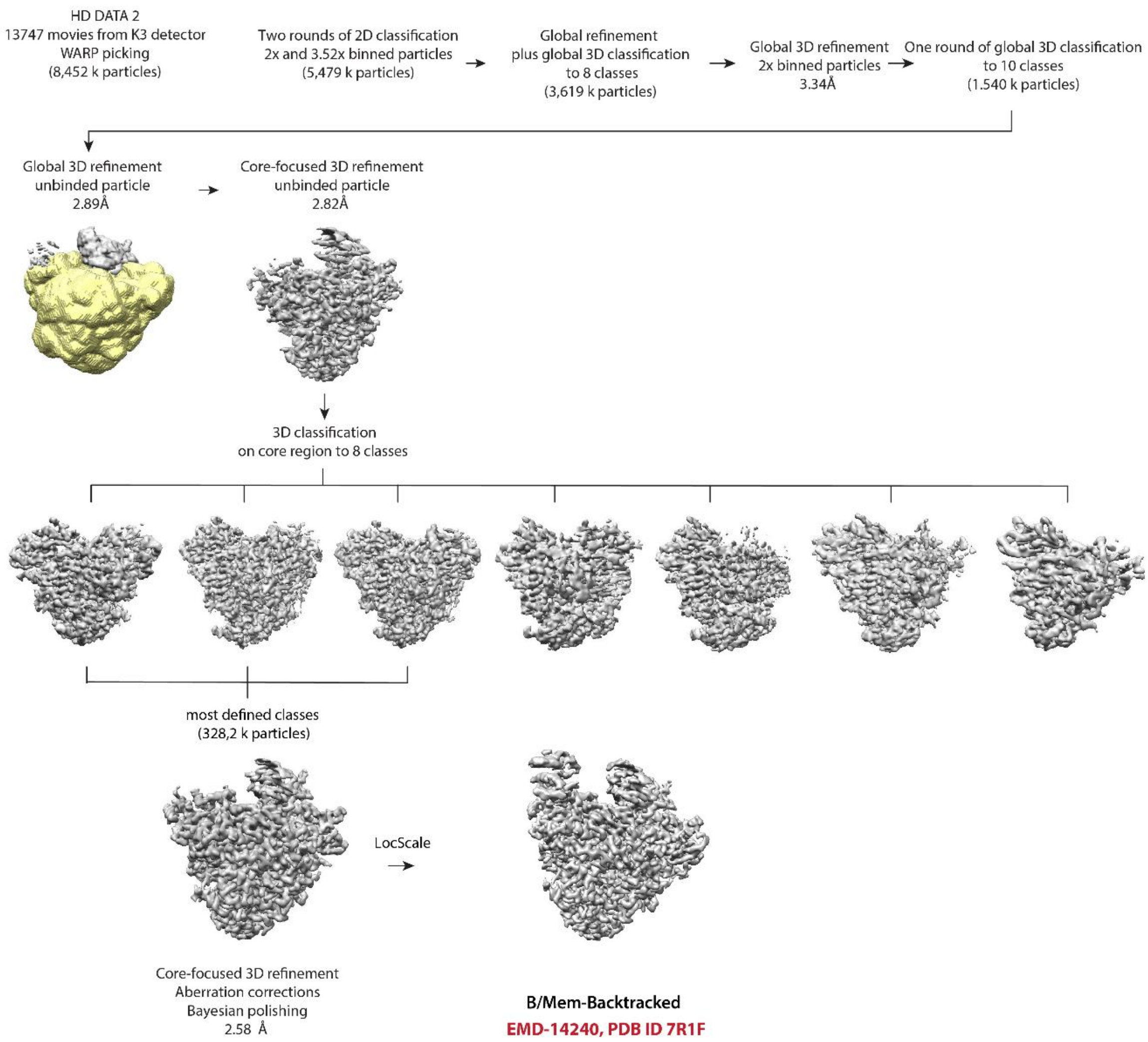


**Figure EM-6: Cryo-EM data 3D classification and refinement scheme of B/Mem-Backtracked complex**

Summary of the cryo-EM 3D classification and refinement scheme of B/Mem-Backtracked complex. HD DATA 2 2D classes with well-defined secondary structure features were merged (5,479 k particles). The merged particles were globally refined and 3D-classified and incomplete, low resolution, and damaged particle classes were excluded from further data analyses. The remaining particles (3,619 k) were globally 3D-refined and subsequently 3D-classified into ten classes. The most defined classes were merged (1,540 k particles), re-extracted to native pixel size, globally and focus 3D-refined and finally focus 3D-classified around the core region of the complex with respective mask (yellow). The most defined classes were merged (328.2 k particles) and core-focus 3D-refined using aberration correction schemes. The final cryo-EM map was refined and post-processed with a respective mask in RELION 3.1 (Zivanov et al., 2018; Zivanov et al., 2020) and filtered by LocScale (Jakobi et al., 2017).

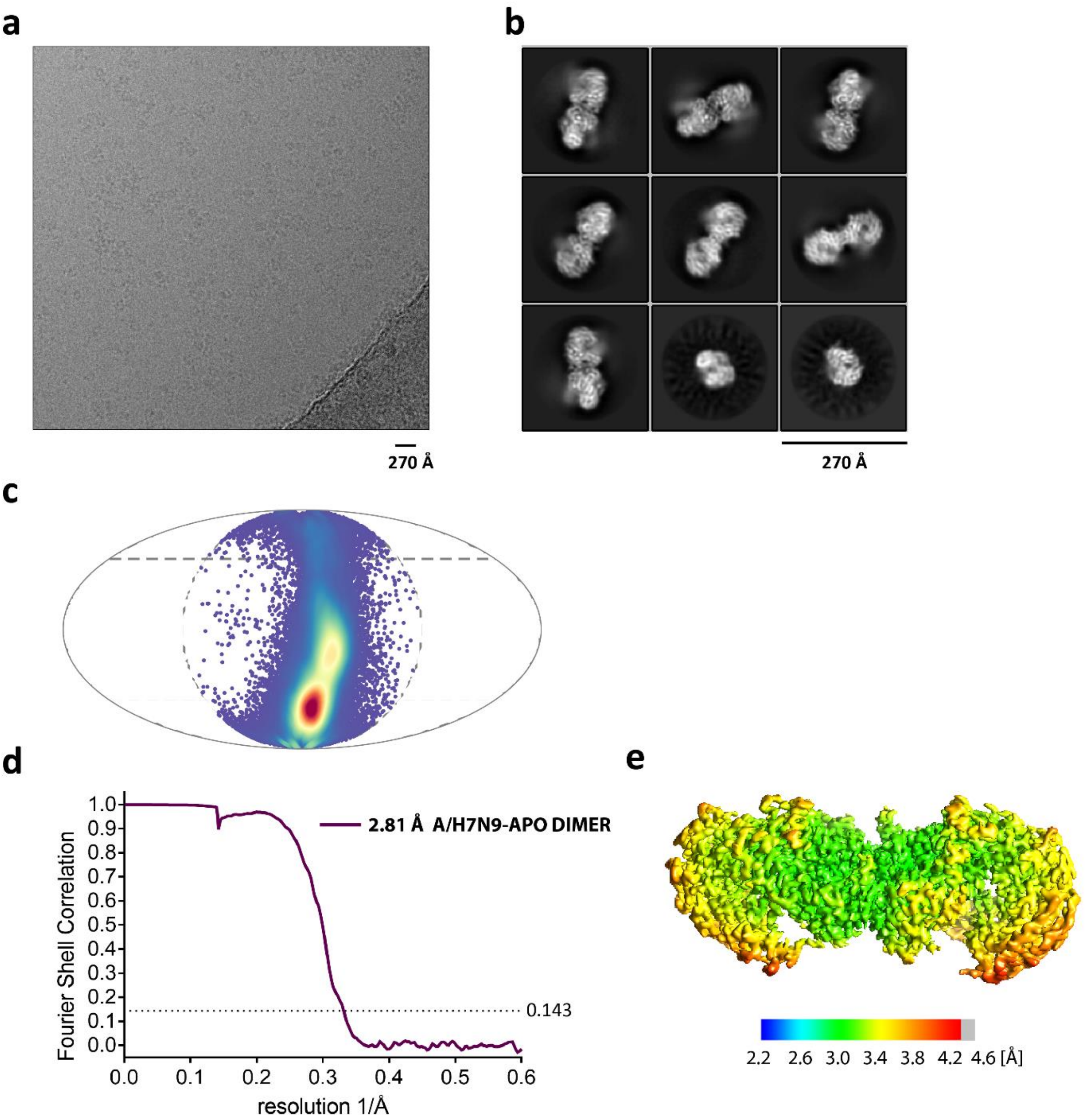


**Figure EM-7: Cryo-EM of A/H7N9-Apo dimer complex**

**a**, Micrograph of the A/H7N9-Apo dimer complex in free standing ice after MotionCor2(Zheng et al., 2017) correction at defocus of ~3 µm. **b**, 2D-class averages of the A/H7N9-Apo dimer complex with minor monomeric fraction **c**, Angular distribution for particle projections in C2 symmetry of the A/H7N9-Apo dimer complex visualized on a globe-like plane. **d**, Fourier shell correlation (FSC) curves for the A/H7N9-Apo dimer complex. The plot of the FSC between two independently refined half-maps shows the overall resolution of the two maps as indicated by the gold standard FSC 0.143 cut-off criteria (Rosenthal and Henderson, 2003). **e**, Surface representation of local resolution distribution of the A/H7N9-Apo dimer complex. The map is colored according to the local resolution (as indicated in the color bar) calculated within the RELION software package.

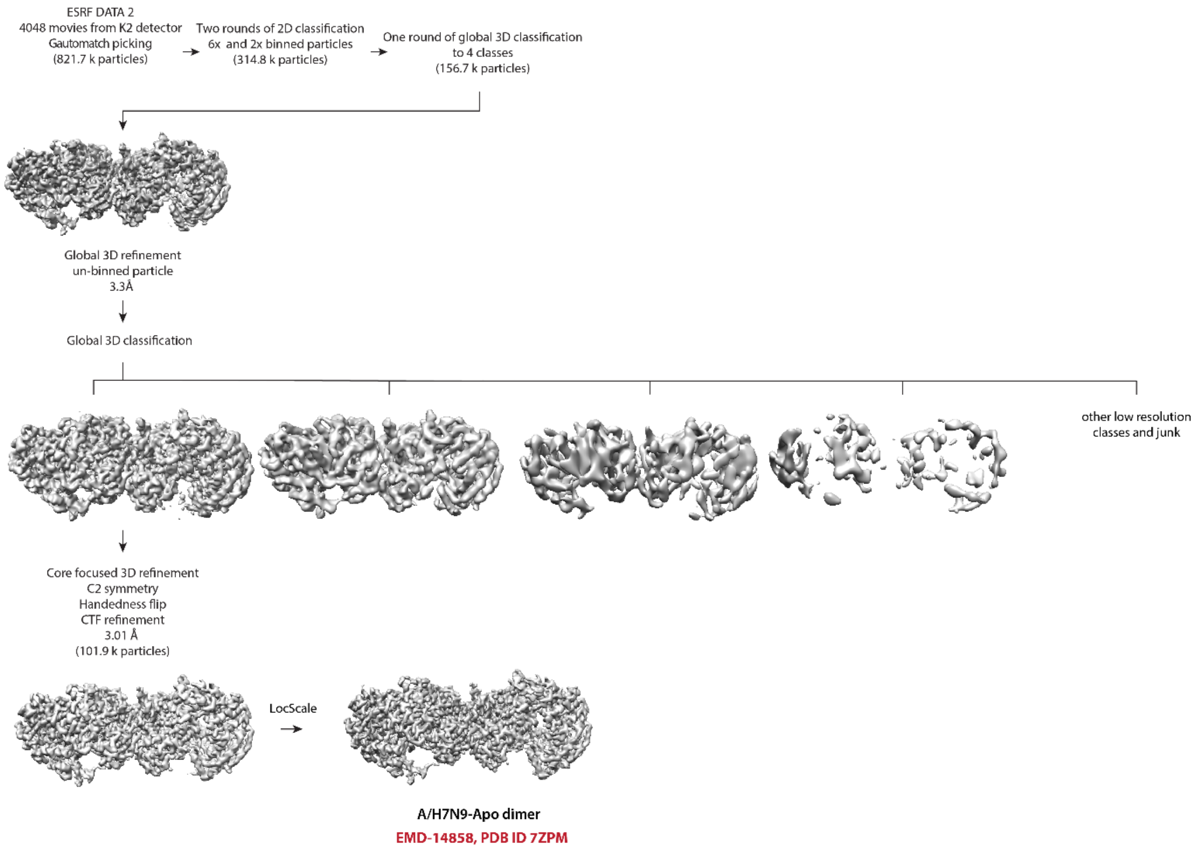


**Figure EM-8: Cryo-EM data 3D classification and refinement scheme of A/H7N9-Apo dimer complex**

Summary of the cryo-EM 3D classification and refinement scheme of A/H7N9-Apo dimer complex. ESRF DATA 2 2D classes with well-defined secondary structure features were merged (314.8 k particles). The merged particles were globally 3D-classified and incomplete, low resolution, and damaged particle classes were excluded from further data analyses. The remaining particles (156.7 k) were globally 3D-refined and subsequently 3D-classified into five classes. The most defined class (101.9 k particles) was globally 3D-refined using C2 symmetry add aberration correction schemes. The final cryo-EM map was refined and post-processed with a respective mask in RELION 3.1 (Zivanov et al., 2018; Zivanov et al., 2020) and filtered by LocScale (Jakobi et al., 2017).

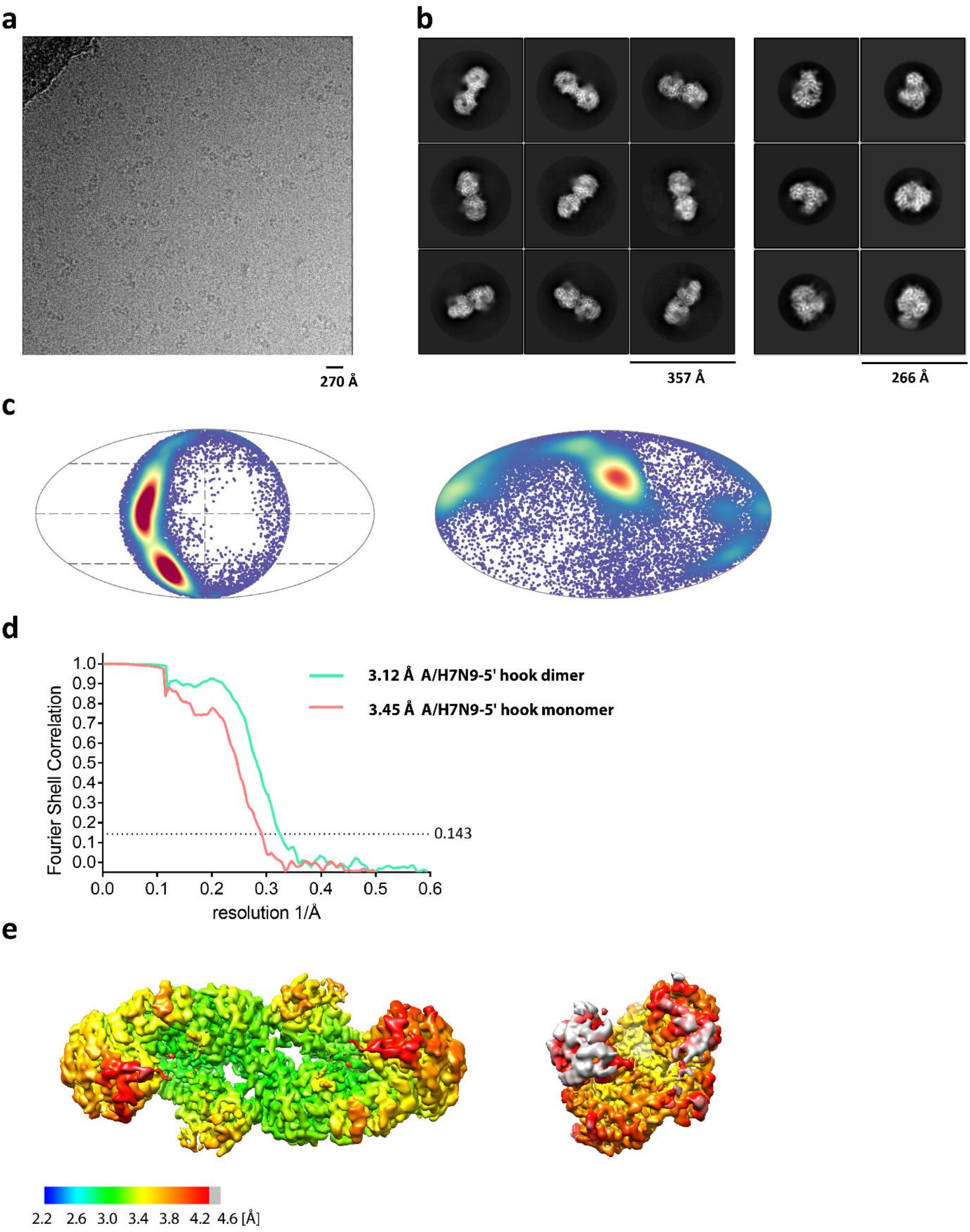


**Figure EM-9: Cryo-EM of A/H7N9-5' hook dimer and monomer complex**

**a**, Micrograph of the A/H7N9-5' hook dimer and monomer complexes in free standing ice after MotionCor2(Zheng et al., 2017) correction at defocus of ~3 µm. **b**, 2D-class averages of the A/H7N9-5' hook dimer (**left**) a monomer (**right**)complex. **c**, Angular distribution for particle projections in C2 symmetry of the A/H7N9-5' hook dimer (**left**) and C1 symmetry of the A/H7N9-5' hook monomer (**right**) complex visualized on a globe-like plane. **d**, Fourier shell correlation (FSC) curves for the A/H7N9-5' hook dimer and monomer complex. The plot of the FSC between two independently refined half-maps shows the overall resolution of the two maps as indicated by the gold standard FSC 0.143 cut-off criteria (Rosenthal and Henderson, 2003). **e**, Surface representation of local resolution distribution of the A/H7N9-5' hook dimer (**left**) and monomer (**right**) complex. The map is colored according to the local resolution (as indicated in the color bar) calculated within the RELION software package.

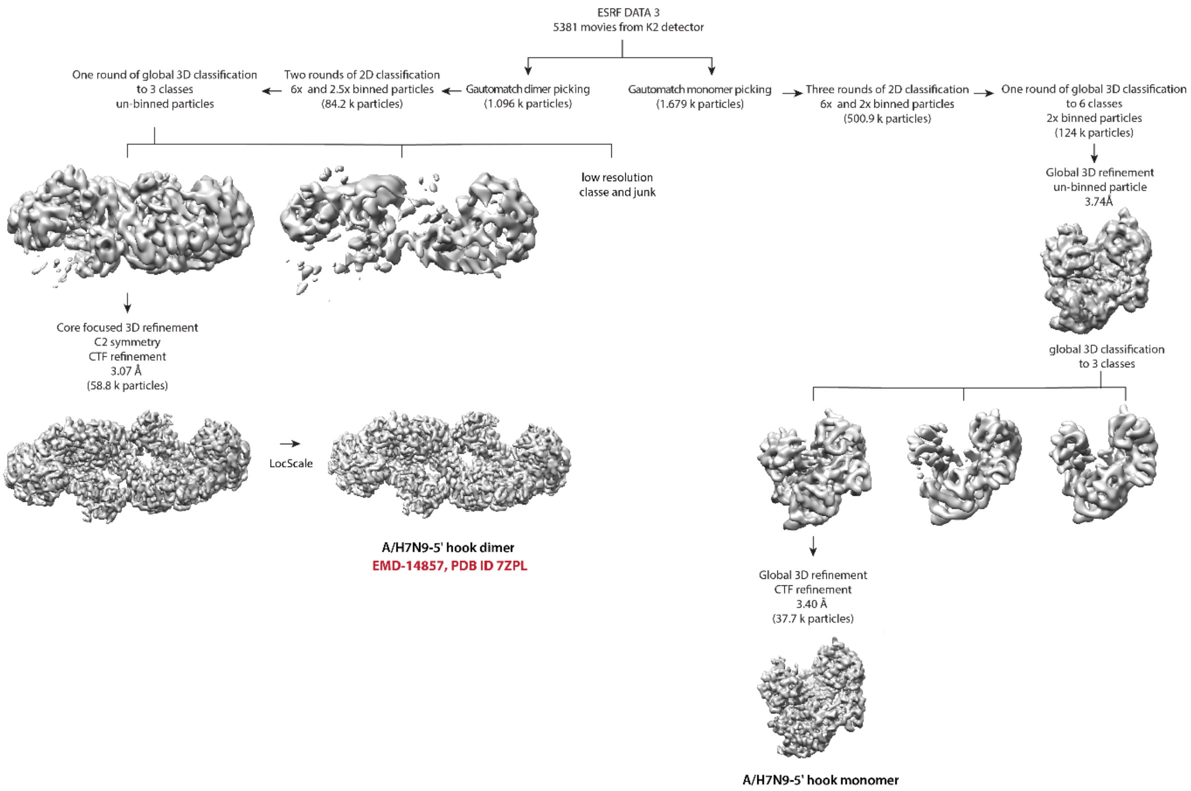


**Figure EM-10: Cryo-EM data 3D classification and refinement scheme of A/H7N9-5' hook dimer and monomer complex**

Summary of the cryo-EM 3D classification and refinement scheme of A/H7N9-5' hook dimer (**left**) and monomer (**right**) complex. Dimeric and monomeric particles were picked using template based picking in Gautomatch and 2D classes with well-defined secondary structure features were merged. The merged A/H7N9-5' hook dimer particles were globally 3D-classified into three classes and the most defined class (58.8 k) was globally 3D-refined using C2 symmetry add aberration correction schemes. The merged A/H7N9-5' hook monomer particles were globally 3D-classified and incomplete, low resolution, and damaged particle classes were excluded from further data analyses. The remaining particles (124 k) were globally 3D-refined and subsequently 3D-classified into three classes. The most defined class (37.7 k particles) was globally 3D-refined using aberration correction schemes. The final cryo-EM map was refined and post-processed with a respective mask in RELION 3.1 (Zivanov et al., 2018; Zivanov et al., 2020) and filtered by LocScale (Jakobi et al., 2017).
